# Lack of evidence for functional luteinising hormone chorionic gonadotropin receptors in non-pregnant human endometrial stromal cells

**DOI:** 10.1101/2022.01.05.474837

**Authors:** ON Mann, C-S Kong, ES Lucas, JJ Brosens, AC Hanyaloglu, PJ Brighton

## Abstract

The human luteinising hormone chorionic gonadotropin receptor (LHCGR) is a G-protein coupled receptor activated by both human chorionic gonadotropin (hCG) and luteinizing hormone (LH), two structurally related gonadotropins with essential roles in ovulation and maintenance of the corpus luteum. LHCGR expression predominates in ovarian tissues where it elicits functional responses through cyclic adenosine mononucleotide (cAMP), Ca^2+^ and extracellular signal-regulated kinase (ERK) signalling. LHCGR has also been localized to the human endometrium, with purported roles in decidualization and implantation. However, these observations are contentious. In this investigation, transcripts encoding *LHCGR* were undetectable in bulk RNA sequencing datasets from whole cycling endometrial tissue and cultured human endometrial stromal cells (EnSC). However, analysis of single-cell RNA sequencing data revealed cell-to-cell transcriptional heterogeneity and identified a small subpopulation of stromal cells with discernible *LHCGR* transcripts. In HEK-293 cells expressing recombinant LHCGR, both hCG and LH elicited robust cAMP, Ca^2+^ and ERK signals that were absent in wild type HEK-293 cells. However, none of these responses were recapitulated in primary EnSC cultures. In addition, proliferation, viability and decidual transformation of EnSC were refractory to both hCG and LH, irrespective of treatment to induce differentiation. Although we challenge the assertion that LHCGR is expressed at a functionally active level in the human endometrium, the discovery of a discrete subpopulation of EnSC that express *LHCGR* transcripts may plausibly account for the conflicting evidence in the literature.

## Introduction

Human chorionic gonadotropin (hCG) and luteinising hormone (LH), like all other gonadotropin hormones, are heterodimers consisting of two monomeric glycoproteins that combine to make a functional protein. They share a common α-subunit, with structural and functional specificity conferred through differing β-subunits. Despite structural similarities, hCG and LH exhibit different temporal profiles and distinct physiological roles (1). In women, LH is produced and released by gonadotropic cells of the anterior pituitary gland, and surges during the mid-luteal phase of the menstrual cycle to drive ovulation and initiate the development of the corpus luteum, a temporary ovarian endocrine structure that produces progesterone. hCG, however, is secreted by the trophectoderm cells of early embryos and eventually the placenta and plays a vital role in sustaining circulating progesterone during early pregnancy.

Within the cycling endometrium, signals from the post-ovulatory rise in progesterone converge with local cyclic adenosine monophosphate (cAMP) production to drive a process of differentiation termed decidualization. This transforms endometrial stromal cells (EnSC) into specialized secretory decidual cells and marks the remodelling of endometrial tissue into the decidua of pregnancy, an immune privileged and nutritive matrix that supports embryo implantation and development (2). However, reprogrammed EnSC can also emerge as acute progesterone-independent senescent decidual cells which propagate within the tissue via inflammatory paracrine secretions and secondary senescence (3, 4). Thus, as progesterone levels fall in the late luteal phase of a non-conception cycle, the decidual tissue is consumed by sterile inflammation and cellular senescence before eventual breakdown and shedding through menstruation (3–5). However, progesterone-dependent decidual cells secrete chemo-attractants to recruit and activate uterine natural killer (uNK) cells, the dominant leukocyte population in the endometrium, which then target and eliminate senescent cells through exocytosis to limit the propagation of senescence (3, 4). Thus, sustained progesterone signalling is essential in maintaining the decidua of pregnancy, and it is embryo-derived hCG that prevents atrophy of the corpus luteum to maintain ovarian progesterone production. This therefore represents an inflection point in the menstrual cycle whereby the fate of the endometrium is principally dependent upon the selection and implantation of embryos that are capable of secreting sufficient hCG and other fitness hormones (6–9).

Both hCG and LH act through the luteinising hormone chorionic gonadotrophin receptor (LHCGR), a 7-transmembrane G-protein coupled receptor (GPCR) comprising a large extracellular domain that binds hCG and LH (1, 10, 11). Activate LHCGR couples to Gα_s_ G-proteins to initiate the adenylyl cyclase, cAMP and protein kinase A (PKA) signalling pathway (1, 10), but it can also activate extracellular signal-regulated kinase (ERK) and protein kinase B (PKB/AKT) to drive cell proliferation, differentiation and survival (1, 12). In addition, at high hormone and receptor levels, LHCGR can also couple to Gα_q_ G-proteins to activate phospholipase C and inositol phosphate signalling to increase intracellular Ca^2+^ (13, 14).

Congruent with its role in ovulation and corpus luteum rescue, LHCGR is prominently expressed in the ovaries (10), but several studies have also demonstrated extra-gonadal expression in tissues of the lower reproductive tract, including the endometrium (15–17). However, these observations have been challenged (18, 19). Several investigations, including papers from the authors, point to endometrial expression, but there are competing claims as to the exact function of the receptors in this tissue (20–29). For example, there is opposing evidence for hCG-mediated generation of intracellular cAMP (21, 25, 26, 30–32), for the role of LHCGR in decidual prolactin production (30, 32–34), and whether expression and function are phasic and cycle-dependent (20, 22, 24, 25, 32).

In the present study, we mined publicly available RNA sequencing data, and despite an absence in bulk sequencing datasets from non-pregnant endometrial tissue and cultured EnSC, we identified a discrete population of stromal cells from single-cell data with detectable *LHCGR* transcripts. However, neither hCG nor LH activated signalling pathways or elicited functional responses in EnSC. Although we found no evidence for functional LHCGR expression in EnSC, heterogeneity in cells expressing the receptor may account for conflicting results between studies.

## Methods

### *in silico* analysis

Receptor expression data were derived from *in silico* analysis of RNA sequencing data within the Gene Expression Omnibus (GEO) repository. Expression from bulk RNA-sequencing within secretory phase endometrial biopsies was derived from GEO accession number GSE65102 (35), and in undifferentiated and decidualized EnSC from GSE104721 (36). Expression within individual cells was analysed from single-cell RNA-sequencing (scRNA-seq) data obtained from whole endometrial biopsies collected 8 and 10 days following the pre-ovulation LH surge, and from EnSC cultures decidualized for 8 days (GEO accession number GSE127918) (3). Single cell expression within endometrial tissues was also derived from an independent data set (37), which was visualized using Cellxgene, a web-based interface for viewing single-cell transcriptomic data (38).

### Endometrial biopsy collection

*Ethical approval and consent:* Endometrial biopsies were obtained from women attending the Implantation Research Clinic at the University Hospitals Coventry and Warwickshire National Health Service Trust. The collection of endometrial biopsies for research was approved by the NHS National Research Ethics - Hammersmith and Queen Charlotte’s & Chelsea Research Ethics Committee (REC reference: 1997/5065) and Tommy’s National Reproductive Health Biobank (REC reference: 18/WA/0356). Written informed consent was obtained prior to tissue collection in accordance with the Declaration of Helsinki, 2000. *Endometrial sampling*: Endometrial biopsies were collected 6-10 days after the pre-ovulatory LH surge using a Wallach Endocell™ Endometrial Cell Sampler and processed immediately. A total of 40 biopsies were used for this investigation and patient demographics are shown in Table S1. *EnSC dissociation and culture*: EnSC were isolated from fresh biopsies using methods described previously (39). In summary, cells were dissociated by mincing and enzymatic digestion of the extracellular matrix using 500 µg/ml collagenase type 1a (Merck Life Sciences UK Ltd, Gillingham, U.K) and 100 µg/ml DNAse I (Lorne Laboratories Ltd, Reading, U.K) for 1 hr at 37°C. Digested tissue was passed through a 40 µM cell strainer to remove glandular clumps and debris and collected via centrifugation (400 *× g*, 5 min). Pellets were re-suspended and cultured in DMEM/F12 media supplemented with 10% dextran-coated charcoal stripped foetal bovine serum (DCC-FBS), antibiotic-antimycotic solution (10 U/ml penicillin, 10 µg/ml streptomycin and 0.25 µg/ml Amphotericin B (Thermo Fisher Scientific, Loughborough, U.K)), 2 mM L-glutamine, 1 nM E_2_ and 2 mg/ml insulin at 37°C in a 5% CO_2_, humidified environment. Growth media was refreshed within 18 hrs to remove non-adherent cells, including immune cells, red blood cells and cellular debris. uNK cells were isolated from this fraction as required (see below). For passage and seeding, EnSC were lifted with 0.25% trypsin, and all treatment protocols were performed in phenol-free DMEM/F-12 media culture containing 2% DCC-FBS and antibiotics/antimycotics. *Cell treatment*: To induce differentiation, EnSC were decidualized with 0.5 mM 8-bromo-cAMP (a cyclic AMP analogue) and 1 µM medroxyprogesterone acetate (MPA) (C+M treatment) as required for individual experiments. Cell were treated with hCG and LH using concentrations and protocols required for individual experiments. Urinary hCG was obtained from Merck Life Sciences UK Ltd (#CG10), and LH from A.F. Parlow, at the National Hormone and Peptide Program, Harbor-UCLA Medical Center).

### HEK-293 cell culture

FLAG-human LHCGR plasmids were constructed and stably transfected into Human Embryonic Kidney (HEK)-293 cells as described previously (40). HEK-LHCGR cells were maintained in DMEM supplemented with 10% DCC-FBS, 1% penicillin/streptomycin and 400 µg/ml Geneticin at 37°C in a 5% CO_2_, humidified environment. Geneticin was excluded from wild-type HEK (HEK-WT) cells. HEK cells were passaged using 0.05% trypsin (< 1 min) and seeded into wells pre-coated with 50 µg/ml poly-D-lysine to aid adherence. Prior to assay, cells were downregulated in serum-free media for 18-24 hrs.

### cAMP assay

Changes in intracellular cellular cAMP were determined using a modified 2-step High-Throughput Time Resolved Fluorescence (HTRF) cAMP kit (Cisbio Bioassay, Codolet, France). Confluent HEK or EnSC cultures in 6-well plates were down-regulated prior to stimulation and assayed in Krebs-Henseleit Buffer (KHB) (composition: NaCl: 118 mM, d-glucose: 11.7 mM, MgSO_4_·7H_2_O: 1.2 mM; KH_2_PO_4_ 1.2 mM, KCl 4.7 mM, HEPES 10 mM, CaCl_2_·2H_2_O: 1.3 mM, pH 7.4), containing 300 µM IBMX. Cells were challenged with hCG or LH (or appropriate controls) for 5 min at 37°C and lysed by rapid aspiration of buffer and immediate addition of 50 µl ice-cold lysis buffer. Cell lysates were collected from wells and cAMP determined on a PHERAstar FS plate reader (BMG Labtech Ltd, Ortenberg, Germany) as per manufacturer’s instructions.

### Ca^2+^ assay

HEK or EnSC cultures were seeded onto 35 mm glass-bottomed culture dishes (MatTek Corporation, MA, U.SA.) and grown until ∼80% confluent. Cells were down-regulated overnight, washed in KHB and loaded with 10 µM Calbryte™ 520 AM (Cal-520) (Stratech Scientific Ltd, Ely, U.K.), a Ca^2+^ sensitive fluorescent dye, for 1 hr at 37°C. Cal-520 was removed prior to assay for 15 min to allow de-esterification of AM esters. Dishes were mounted onto the stage of an Olympus IX-85 microscope and equilibrated at 37°C in KHB before imaging. Cells were challenged with 1 µM hCG or LH and Cal-520 fluorescence captured by a DC4100 LED stack and GFP filter set (ThorLabs, Newton, NJ, U.S.A.). Images were obtained using a Zyla sCMOS camera (Andor Technology Ltd, Belfast, U.K.) at a frequency of 1 Hz. Videos were analysed in ImageJ and changes in cytosolic fluorescence relative to time 0 (F/F_0_) were used as an index of intracellular Ca^2+^. For mean data, peak responses were obtained from 12 cells chosen at random from each independent primary culture.

### Western blot

EnSC and HEK cultures were grown to ∼80% confluency in 6-well plates and down-regulated for 24 hrs prior to assay. Where required, EnSC were decidualized for 8 days prior to assay. Cells were assayed in KHB at 37°C and challenged with 10 nM hCG or LH for up to 45 min. Experiments were terminated by aspiration of buffer and rapid lysis in ice-cold RIPA buffer containing cOmplete mini protease inhibitors (Roche, Basel, Switzerland) and phosphatase inhibitor cocktail 2 (1:1000) (Merck Life Sciences U.K. Ltd). Cell debris was removed from lysates via centrifugation (275 × *g*, 2 min, 4°C), and samples were prepared in 25% (v:v) NuPage LDS x4 sample buffer (Fisher Scientific, Loughborough, U.K.) and 100 nM DTT (Merck Life Sciences U.K. Ltd), and heated at 95°C for 5 min. Proteins were separated using standard SDS-PAGE electrophoresis techniques and transferred onto 0.45 μm nitrocellulose membranes (GE Healthcare, Amersham, U.K.) where non-specific binding was blocked by 5% (w:v) powdered milk in TBS-T (50 mM Tris, 150 mM NaCl, pH 7.4, 0.5% (v:v) Tween-20) for 1 hr at RT. Membranes were probed with antibodies targeting phosphorylated-ERK (1:2000 in TBS-T, Cell Signalling Technologies, MA, U.S.A, antibody #9101) overnight at 4°C. Blots were washed clear of unbound antibody in TBS-T before addition of anti-rabbit-HRP secondary antibody (1:1000 in 5% milk/TBS-T; Agilent Technologies, CA, U.S.A.) for 1 hr at RT. Blots were washed and visualized using ECL reagent (GE Healthcare) and auto-radiography. Protein loading was controlled by uniform cell seeding but also verified by detection of total-ERK. Here, antibodies were stripped from membranes by submersion in boiling water for 5 min. and. Membranes were blocked in 5% milk/TBS-T and probed with antibodies targeting total-ERK (1:2000 in TBS-T, Cell Signalling Technologies, antibody #9102). The density of phospho-ERK bands was determined using GeneTools gel analysis software (Syngene, Cambridge, U.K.), and expressed relative to levels of total-ERK.

### XTT cell viability Assay

EnSC were grown in 96-well plates and down-regulated for 24 hrs prior to a 2-day treatment with hCG or LH, both with and without C+M treatment to induce decidualization. The viability of EnSC after treatment was assessed using the CyQuant™ XTT cell viability assay (Fisher Scientific) as per manufacturer’s instructions.

### uNK cell isolation and counting

uNK cells were isolated and cultured as described elsewhere (4, 39). Briefly, unattached cells within culture supernatant from 3-4 freshly digested endometrial biopsies were pooled following overnight culture. Red blood cells were depleted via Ficoll-Paque density medium, and suspensions were incubated with magnetic microbeads targeting the uNK specific antigen CD56 (#130-050-401, Miltenyi Biotec, Bergisch Gladbach, Germany) diluted 1:20 in wash buffer (1% BSA/PBS) for 15 min at 4°C. CD56^+^ cells were isolated through magnetic activated cell sorting (MACS) and cultured in DMEM/F12 media supplemented with 10% DCC-FBS, 1% antibiotic/antimycotic and 125 pg/mL recombinant IL-15. Isolated uNK cells were treated with 10 nM hCG and viability and proliferation were quantified by XTT assay and cell counting using a Neubauer-improved haemocytometer after 2 days.

### xCELLigence

Cell proliferation was assessed in real-time using an xCELLigence^®^ Real Time Cell Analysis (RTCA) DP instrument (ACEA Biosciences Inc, San Diego, CA, U.S.A). EnSC were seeded in E-16 xCELLigence plates at a density of 10,000 cells/well and cultured for 24 hrs. Cells were down-regulated in 2% DCC-FBS overnight prior to assay. Both undifferentiated and decidualizing cells (C+M treated) were monitored in the presence of hCG and LH with electrical impedance referenced to 0 before treatment. Changes in impedance were captured and analysed using the RTCA Software v1.2 to reflect changes in cell adherence, shape and proliferation and were expressed as the arbitrary unit ‘cell index’.

### Senescence Associated-β-Galactosidase

Levels of the cellular senescence marker Senescence Associated-β-Galactosidase (SA-β-Gal) were determined using a modified version of the Quantitative Cellular Senescence Assay kit (Cell Biolabs Inc; CA, U.S.A.). Both undifferentiated and decidualized (C+M treated) EnSC in 96-well plates were treated with 10 nM hCG or LH for 8 days. Cells were washed in ice-cold PBS and lysed in 50 μL ice-cold assay lysis buffer containing cOmplete™ mini protease inhibitors. Lysates (25 µL) were transferred to black-walled, black-bottomed 96-well plates and an equal volume of 2 x assay buffer was added as per manufacturer’s guidelines. Plates were sealed to avoid evaporation and incubated for 1 hr at 37°C in a non-humidified, non-CO_2_ incubator. The reaction was terminated by 200 μL stop solution and fluorescent intensity units (F.I.U.) were determined on a PHERAstar FS plate reader at 360/465 nm. Assays were normalized by identical cell seeding.

### RT-qPCR

Total RNA from EnSC was extracted using STAT-60 (AMSBio, Abingdon, U.K.), according to manufacturer’s instructions, with recovered RNA quantified on a Nanodrop spectrophotometer. Equal amounts of RNA were transcribed into cDNA using the QuantiTect Reverse Transcription Kit (Qiagen, Manchester, U.K.) as per manufacturer’s instructions. Target gene expression was analysed using Quantifast SYBR Green Master Mix (Fisher Scientific) on a 7500 Fast Real-Time PCR System (Applied Biosystems, CA, U.SA.). The expression level of each gene was calculated using the ΔCt method and normalized against *L19* housekeeping gene. Primer sequences were: *PRL* sense 5’-AAG CTG TAG AGA TTG AGG AGC AAA C-3’, *PRL* antisense 5’-TCA GGA TGA ACC TGG CTG ACT A-3’, *IGFBP1* sense 5’-CGA AGG CTC TCC ATG TCA CCA-3’, *IGFBP1* antisense 5’-TGT CTC CTG TGC CTT GGC TAA AC-3’, *IL1RL1* sense 5’-TTG TCC TAC CAT TGA CCT CTA CAA-3’, *IL1RL1* antisense 5’-GAT CCT TGA AGA GCC TGA CAA-3’, *CLU* sense 5’-GGG ACC AGA CGG TCT CAG-3’, *CLU* antisense 5’-CGT ACT TAC TTC CCT GAT TGG AC-3’; *L19* sense 5’-GCG GAA GGG TAC AGC CAA-3’, *L19* antisense 5’-GCA GCC GGC GCA AA-3’

### ELISA

Supernatant from cell cultures was collected every 2 days and cleared of cellular debris by centrifugation (275 × *g*, 5 min, RT) before storage at -20°C. Levels of secreted IGFBP1, prolactin, clusterin and soluble ST2 (sST2) in culture supernatant were quantified using DuoSet ELISA kits (Biotechne, MN, U.S.A.) as per manufacturer’s instructions. Absorbance at 450 nm was determined on a PHERAstar FS plate reader with concentrations interpolated from known standards using a 4-parameter fit in GraphPad Prism (v9.0).

### Statistical analysis

GraphPad Prism v9.0 (GraphPad Software Inc. CA, U.S.A.) was used for statistical analyses with data presented as fold-change relative to the most informative comparator. Paired Student’s *t*-test was performed to determine statistical significance between 2 groups, and one-way ANOVA, followed by Tukey’s or Dunnett’s post-hoc test for multiple comparisons involving more than 2 groups. Data with 2 independent variables was analysed using a two-way ANOVA (mixed methods) and Dunnett’s multiple comparison test using the most appropriate control as the comparator. In all cases, *P*<0.05 was considered significant.

## Results

### Lack of *LHCGR* transcripts in the endometrium

A summary of the literature pertaining LHCGR expression in endometrial cells is shown in Table 1 and highlights contradictory evidence on this subject. We therefore explored LHCGR expression within the endometrium by mining publicly available RNA-sequencing (RNA-seq) data from whole endometrial biopsies and EnSC cultures (GEO accession numbers: GSE65102 and GSE104721). Within whole tissue, transcripts encoding *LHCGR* were undetectable in 8 out of 20 biopsies. Average levels were 0.15 ± 0.26 transcripts per million (TPM) (mean ± SD) and significantly lower (*P*<0.05) than those encoding other GPCRs implicated in endometrial regulation, including the PGE_2_ receptor (*PTGER2*), relaxin receptor (*RXFP1*) and the lysophosphatidic acid receptor (*LPAR1*) (Fig. 1A). Equally, in EnSC, *LHCGR* transcripts were absent in 2 out of 3 cultures, whereas those encoding other GPCRs including *PTGER2*, *RXFP1*, *LPAR1* and the oxytocin receptor (*OXTR*), were prominent (Fig. 1B). One disadvantage of bulk RNA-sequencing is that it masks subpopulation variability and heterogeneity between cells. Single-cell RNA sequencing (scRNA-seq) however, enables molecular characterisation of individual cells, and is a powerful tool to overcome these limitations by revealing transcriptional heterogeneity within cell populations (41). We therefore mined scRNA-seq data that reconstructed the decidual pathway by treating EnSC with 8-bromo-cAMP and the progestin medroxyprogesterone (MPA) (C+M treatment) over 8 days (GEO accession number: GSE127918). Cells clustered into distinct cell states based on their transcriptional profile which closely mirrored the treatment regime (Fig. S1A). Irrespective of cell state, the average expression of *LHCGR* transcripts was low when compared to *RXFP1*, *LPAR1* and *OXTR*. However, one striking observation was the considerable heterogeneity within the EnSC population for all receptors, including LHCGR (Fig. 1C). Out of the 4574 cells in the dataset we identified 97 (2.1%) cells where transcripts encoding *LHCGR* were detected (Fig. 1D).

**Figure 1.**
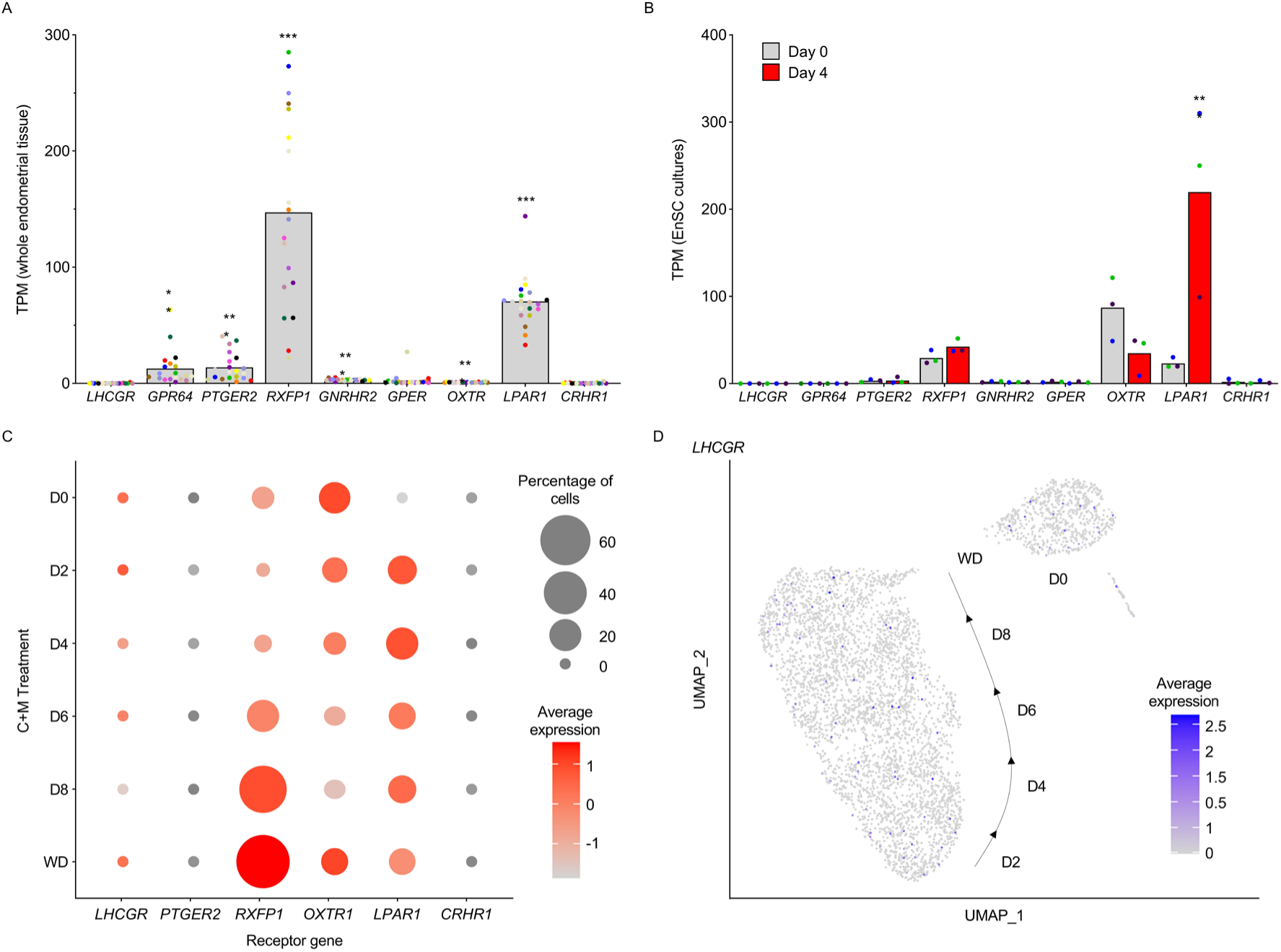
LHCGR expression in whole endometrial biopsies and EnSC. A) Expression of genes coding GPCRs within whole endometrial biopsies (GEO accession no. GSE65102). Data points from individual patients are colour-matched and shown together with bar graphs denoting mean (n= 20). Significance was determined by ANOVA and Dunnett’s multiple comparison test using *LHCGR* as the comparator. ** denotes *P*<0.01 and *** *P*<0.001. B) Expression of GPCRs in undifferentiated EnSC (day 0) and cells decidualized with 8-br-cAMP and MPA (C+M) for 4 days (GEO GSE104721). Individual data points from 3 biological replicates are shown with bars denoting mean. Significance was determined by two-way ANOVA and Sidak’s multiple comparison test, *** denotes *P*<0.001. C) Dot-plot to visualize scRNA-seq data (GEO GSE127918) showing the percentage of EnSC expressing GPCRs, and the Log_2_-transformed average expression within these cells. D) UMAP (uniform manifold approximation and projection) projections of *LHCGR* expression within scRNA-seq data from undifferentiated (D0) EnSC and those treated with C+M for time points indicated before withdrawal (WD). Transcriptomic profiling to identify cell states is shown in Fig. S1A.

**Table 1.**
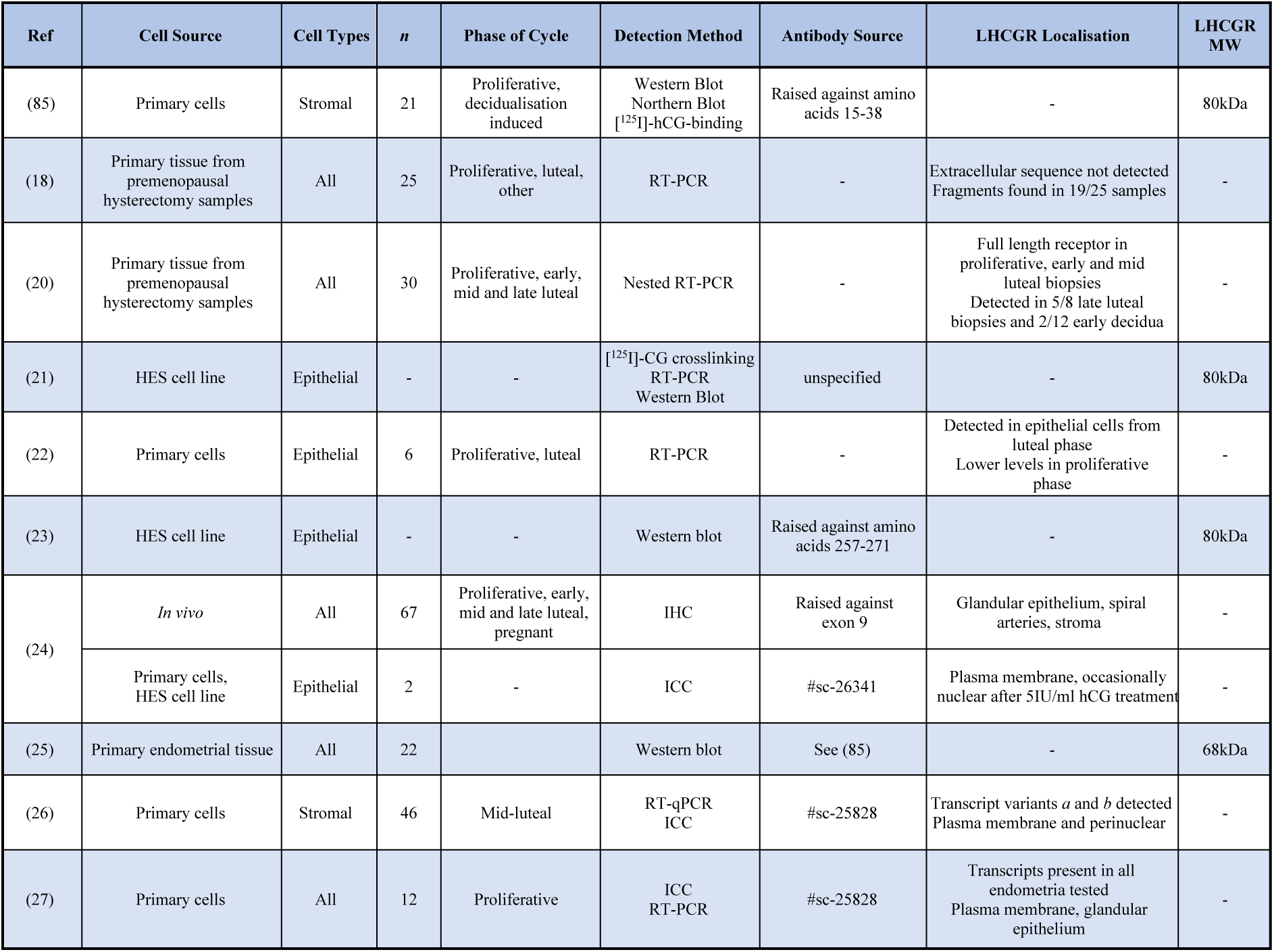
Expression of LHCGR in endometrial cells and tissue

| Ref | Cell Source | Cell Types | *n* | Phase of Cycle | Detection Method | Antibody Source | LHCGR Localisation | LHCGR MW |
| --- | --- | --- | --- | --- | --- | --- | --- | --- |
| (85) | Primary cells | Stromal | 21 | Proliferative, decidualisation induced | Western Blot<br>Northern Blot<br>[<sup>125</sup>I]-hCG-binding | Raised against amino acids 15-38 | - | 80kDa |
| (18) | Primary tissue from premenopausal hysterectomy samples | All | 25 | Proliferative, luteal, other | RT-PCR | - | Extracellular sequence not detected<br>Fragments found in 19/25 samples | - |
| (20) | Primary tissue from premenopausal hysterectomy samples | All | 30 | Proliferative, early, mid and late luteal | Nested RT-PCR | - | Full length receptor in proliferative, early and mid luteal biopsies<br>Detected in 5/8 late luteal biopsies and 2/12 early decidua | - |
| (21) | HES cell line | Epithelial | - | - | [<sup>125</sup>I]-CG crosslinking<br>RT-PCR<br>Western Blot | unspecified | - | 80kDa |
| (22) | Primary cells | Epithelial | 6 | Proliferative, luteal | RT-PCR | - | Detected in epithelial cells from luteal phase<br>Lower levels in proliferative phase | - |
| (23) | HES cell line | Epithelial | - | - | Western blot | Raised against amino acids 257-271 | - | 80kDa |
| (24) | *In vivo* | All | 67 | Proliferative, early, mid and late luteal, pregnant | IHC | Raised against exon 9 | Glandular epithelium, spiral arteries, stroma | - |
|  | Primary cells, HES cell line | Epithelial | 2 | - | ICC | #sc-26341 | Plasma membrane, occasionally nuclear after 5IU/ml hCG treatment | - |
| (25) | Primary endometrial tissue | All | 22 |  | Western blot | See (85) | - | 68kDa |
| (26) | Primary cells | Stromal | 46 | Mid-luteal | RT-qPCR<br>ICC | #sc-25828 | Transcript variants *a* and *b* detected<br>Plasma membrane and perinuclear | - |
| (27) | Primary cells | All | 12 | Proliferative | ICC<br>RT-PCR | #sc-25828 | Transcripts present in all endometria tested<br>Plasma membrane, glandular epithelium | - |

Additionally, the major cell types within whole endometrial biopsies (GEO accession number: GSE127918) were deconvolved and identified based on their transcriptomic profile (Fig. S1B, left panel), and consistent with EnSC, *LHCGR* transcripts were detected in a subpopulation of stromal and epithelial cells (Fig. S1B, right panel). To further substantiate the heterogeneity in whole tissue we mined independent single cell transcriptomic data (37), and again observed *LHCGR* transcripts in a small population of stromal cells (Fig. S2D-E).

### hCG and LH do not elicit signalling pathways in EnSC

A summary of the literature investigating the signalling and functional responses of primary endometrial cells to hCG and LH is summarized in Table 2, and highlights the opposing evidence and contention in this area.

**Table 2.** Functional activation of LHCGR in primary endometrial cells

| Ref | Cell Source | Cell Types | n | Phase of Cycle | hCG Source | hCG Concentration | LHCGR Response | % change in response (relative to control) |
| --- | --- | --- | --- | --- | --- | --- | --- | --- |
| (86) | Primary cells | Stromal | 27 | Proliferative | National Hormone and Pituitary Program | 1.49-14900 IU/ml | Cox-2 protein levels by Western Blot | 100% increase |
|  |  |  |  |  |  |  | Cox-2 transcript levels by Northern blot | Information not given, only statistically significant above 149 IU/ml |
| (31) | Primary cells | Stromal | 17 | Luteal | Sigma, #994F-0137 | 50, 250, or 500 ng | cAMP concentration by radioimmunoassay | 500% increase |
| (85) | Primary cells | Stromal | 21 | Proliferative | National Hormone and Pituitary Program | 149 IU/ml | LHCGR transcript levels by Northern blot | 60% decrease |
|  |  |  |  |  |  |  | LHCGR expression by Western blot | 40% decrease |
| (28) | Primary cells | Stromal | 20 | Secretory | Sigma | 10 nM | PRL expression via RT-qPCR | 60% decrease in control patients<br>20% increase in RPL patients |
|  |  |  |  |  |  |  | PROK1 expression via RT-qPCR | 50% decrease in control patients<br>350% increase in RPL patients |
| (72) | Primary cells | Stromal | 5 | - | - | 0.1 – 100 IU/ml | PRL via RT-qPCR and ELISA | 58% decrease with in mRNA and protein |
|  |  |  |  |  |  |  | IGFBP1 via RT-qPCR and ELISA | 50% decrease in mRNA and protein |
| (33) | Primary cells | Stromal | 24 | Proliferative | National Hormone and Pituitary Program | 1.49-149 IU/ml | Morphology by phase contrast microscopy | Morphological changes consistent with decidualisation |
|  |  |  |  |  |  |  | PRL protein levels by radioimmunoassay | No change, but 200% increase in combination with E<sub>2</sub>+P<sub>4</sub> |
| (32) | Primary cells | Stromal | 68 | Follicular, periovulatory, early-late luteal | Teikokuzouki pharmaceuticals | 0.01-100 IU/ml | PRL protein levels by radioimmunoassay | 100% decrease at 100 IU/ml. No change at 1, 10 IU/ml |
|  |  |  |  |  |  |  | cAMP concentration by radioimmunoassay | 66% decrease at 100IU/ml. No change at 1, 10 IU/ml |
| (21) | HES cell line | Epithelial | - | - | National Hormone and Pituitary Program | 1.8 IU/ml | cAMP concentration by ELISA | Statistically insignificant |
|  |  |  |  |  |  |  | phospho-ERK by Western blot | Increased band intensity (not quantified) |
|  |  |  |  |  |  |  | PGE<sub>2</sub> production by ELISA | 100% increase |
| (22) | Primary cells | Epithelial | 28 | Proliferative, luteal | Pregnyl, Organon | 1-50 IU/ml | LIF production by ELISA | 100-125% increase |
|  |  |  |  |  |  |  | IL-6 production by ELISA | 20% decrease |
|  |  |  |  |  |  |  | Cell proliferation by BrdU ELISA | Statistically insignificant |
| (87) | Primary cells | Epithelial | 18 | Proliferative, luteal | Pregnyl, Organon, Ovitrelle, Serono | 5-50 IU/ml | VEGF protein levels by ELISA | 60% increase |
| (23) | HES cell line | Epithelial | - | - | EMD Serono | 10 IU/ml | phospho-ERK by Western blot | 400% increase |
| (88) | Primary cells | Epithelial | 15 | Proliferative, luteal | National Hormone and Peptide Program | 0.2-20 IU/ml | FGF2 protein levels, among others, by multiplex immunoassay | 50% increase |
| (24) | Primary cells HES cell line | Epithelial | 2 | - | National Hormone and Pituitary Program | 20 IU/ml (acute) or 0.5-5 + 20 IU/ml (chronic) | phospho-ERK by Western Blot | 40-50% increase |
|  |  |  |  |  |  |  | Transepithelial resistance | 40% decrease (acute dose) |
|  |  |  |  |  |  |  | Cell adhesion | 20-50% increase |
| (25) | Primary cells | All | 22 | Various | Source not specified | 10µg/ml | Adenylyl cyclase activity by radioactive cAMP production | Statistically insignificant |
| (26) | Primary cells | Stromal | 46 | Mid-luteal | Sigma (#CG10) | 10 IU/ml | cAMP concentration by ELISA | 100% increase |
|  |  |  |  |  |  |  | phospho-ERK by Western blot | 100% increase |
| (27) | Primary cells | All | 12 | Proliferative | Luveris, Merck Serono | 0-100 ng/ml | cAMP concentration by ELISA | 200% increase |
|  |  |  |  |  |  |  | CYP191A and P450ssc expression by RT-qPCR | 300% and 200% increase, respectively |

As shown in Fig. 2A, active LHCGR couples to Gα_s_ G-proteins to activate adenylate cyclase and increase intracellular cAMP, but it can also trigger Ca^2+^ release and activate kinase pathways including ERK. To validate gonadotropin bioactivity, these key signalling pathways were investigated in HEK-293 cells recombinantly expressing LHCGR (HEK-LHCGR). Cells were challenged with either hCG or LH and compared to parallel experiments in wild-type HEK-293 cells (HEK-WT). Robust increases in intracellular cAMP were detected in HEK-LHCGR cells challenged for 5 minutes with either hCG or LH. Inductions were concentration-dependent and generated pEC_50_ values of -9.671 (213pM) for hCG and -9.370 (426pM) for LH, and maximal responses were achieved at ∼10 nM (Fig. 2B). By contrast, HEK-WT cells did not respond. HEK cells were also loaded with the Ca^2+^ sensitive dye Calbryte-520 and challenged with supra-maximal concentrations (1 µM) of hCG or LH. Both agonists elicited robust Ca^2+^ responses in HEK-LHCGR cells, but not HEK-WT cells (Fig. 2C). Further, both ligands induced a strong and sustained induction of phosphorylated (active)-ERK in HEK-LHCGR, whereas HEK-WT cells did not respond (Fig 2D).

**Figure 2.**
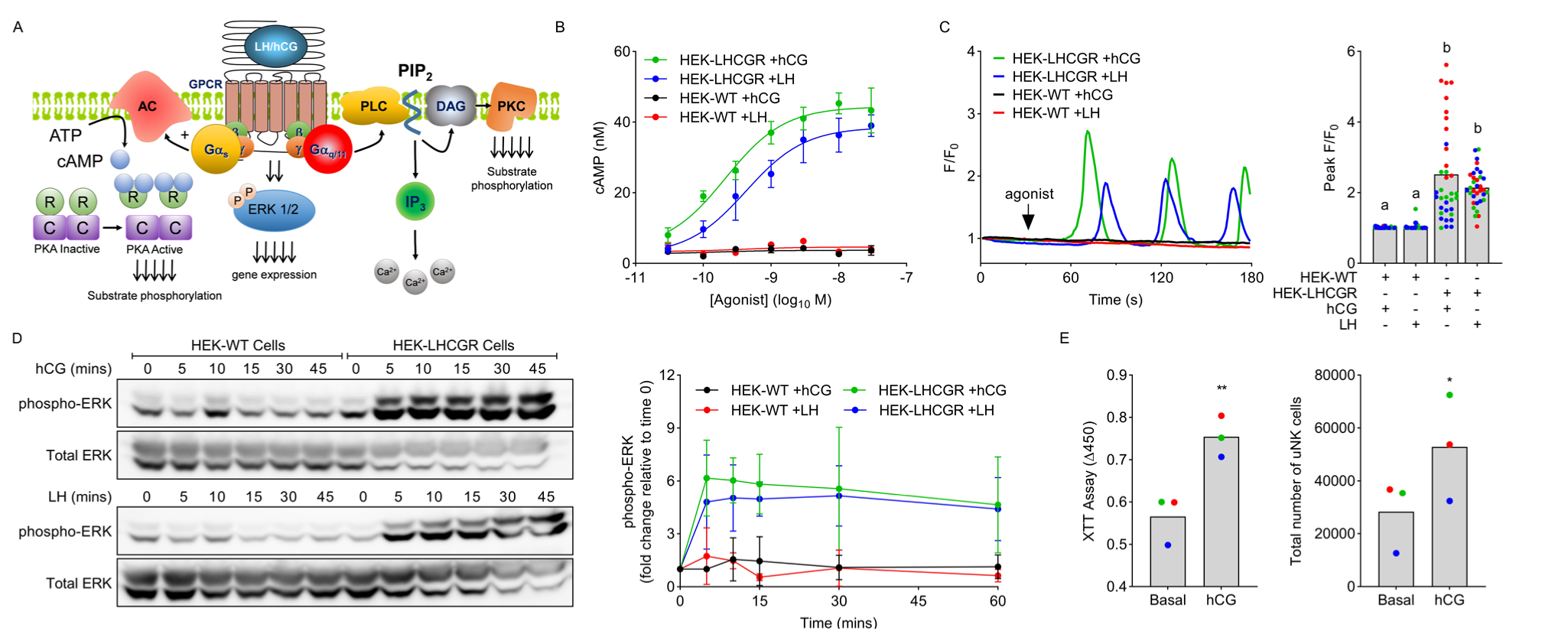
hCG and LH are bioactive. A) Schematic representation of signalling pathways downstream of LHCGR. B) Concentration-dependent cAMP responses to hCG and LH in HEK-WT and HEK-LHCGR cells. C) Representative Ca^2+^ traces in HEK-WT and -LHCGR cells challenged with 1 µM hCG or LH. Data represent one individual cell. Mean peak responses from 36 cells analysed from 3 independent cultures (denoted as separate colours) are shown in the right panel with differing letters indicating significance between groups (*P*<0.05) (ANOVA and Tukey’s multiple comparison test). D) Representative western blots showing levels of phospho-ERK in HEK-WT and -LHCGR cells challenged with either 10 nM hCG (top panel) or LH (lower panel). Mean data ± SD from 3 separate cultures is shown in the right panel. E) Impact of 10 nM hCG on uNK cell proliferation as measured by XTT assay (left panel) and cell counting (right panel). Individual data points from 3 biological replicates are shown with bars denoting mean values. Significance was determined by paired t-test with * = *P*<0.05 and ** *P*<0.01.

In addition to LHCGR, LH and hCG also bind with mannose and other C-type lectin receptors (42–45). Significantly, hCG-mediated mannose receptor activation has been shown to induce uNK cell proliferation (44). To demonstrate alternative hCG-receptor interactions, and to provide further evidence of bioactivity, we investigated the proliferative effects of hCG on uNK cells isolated from non-pregnant endometrial biopsies. We have previously demonstrated that isolated uNK cells are viable, and stain positively for both the pan-leukocyte marker CD45 and the uNK specific marker CD56 (8). In this study, uNK cell proliferation, as quantified by XTT assay, was augmented by 133 ± 35% (*P*=0.0093) (mean ± SD) after 2 days in culture with 10 nM hCG (Fig. 2E, left panel). Equally, the overall number of uNK cells, as determined via haemocytometer, increased by 202 ± 54% (*P*=0.030) when cultured with hCG (Fig. 2E, right panel). These observations prompted us to characterize the expression profiles of mannose receptors within the endometrium. First, we mined publicly available RNA-seq data from endometrial biopsies (GEO accession number: GSE65102) and found that transcripts for both the mannose receptor C-type 1 (*MRC1*) and type 2 (*MRC2*) were abundant, and ∼48 (*P*<0.0001) and ∼443 (*P*<0.0001) fold higher than those for *LHCGR*, respectively (Fig. S2A). Next, we examined expression in EnSC (GEO accession number: GSE104721) and observed high levels of transcripts for *MRC2*, but none for *MRC1* (Fig. S2B). Likewise, *MRC1* transcripts were absent within our whole tissue scRNA-seq data (GEO accession number: GSE127918), whereas *MRC2* was prevalent in the stromal compartment and immune cells, including lymphocytes (Fig. S2C). Similar observations from independent scRNA-seq data also indicated an absence of *MRC1* transcripts in most cell types except a small population of macrophages, but widespread *MRC2* expression that predominated in the stromal compartment and immune cells (Fig. S2D-F). Interestingly, for all receptors there was conspicuous heterogeneity between individual cells.

Having validated the ligands, we next sought to investigate hCG and LH-mediated signalling in EnSC. In contrast to responses in HEK-LHCGR, neither hCG nor LH increased intracellular cAMP in primary EnSC cultures. Cells were however responsive to PGE_2_ (*P*=0.037), known to activate Gα_s_-coupled EP_2_/EP_4_ prostanoid receptors, and forskolin (*P*<0.0001), the receptor-independent adenylate cyclase activator (Fig. 3A). Further, the same supra-maximal concentration (1 µM) of hCG and LH that elicited robust Ca^2+^ responses in HEK-LHCGR cells failed to increase intracellular Ca^2+^ in EnSC, despite strong responses to PGE_2_, likely activating the Gα_q/11_-coupled EP_1_/EP_3_ prostanoid receptors (Fig. 3B). Likewise, basal levels of phosphorylated-ERK in both undifferentiated EnSC and those decidualized for 8 days were unchanged following stimulation with either hCG or LH (Fig. 3C).

**Figure 3.**
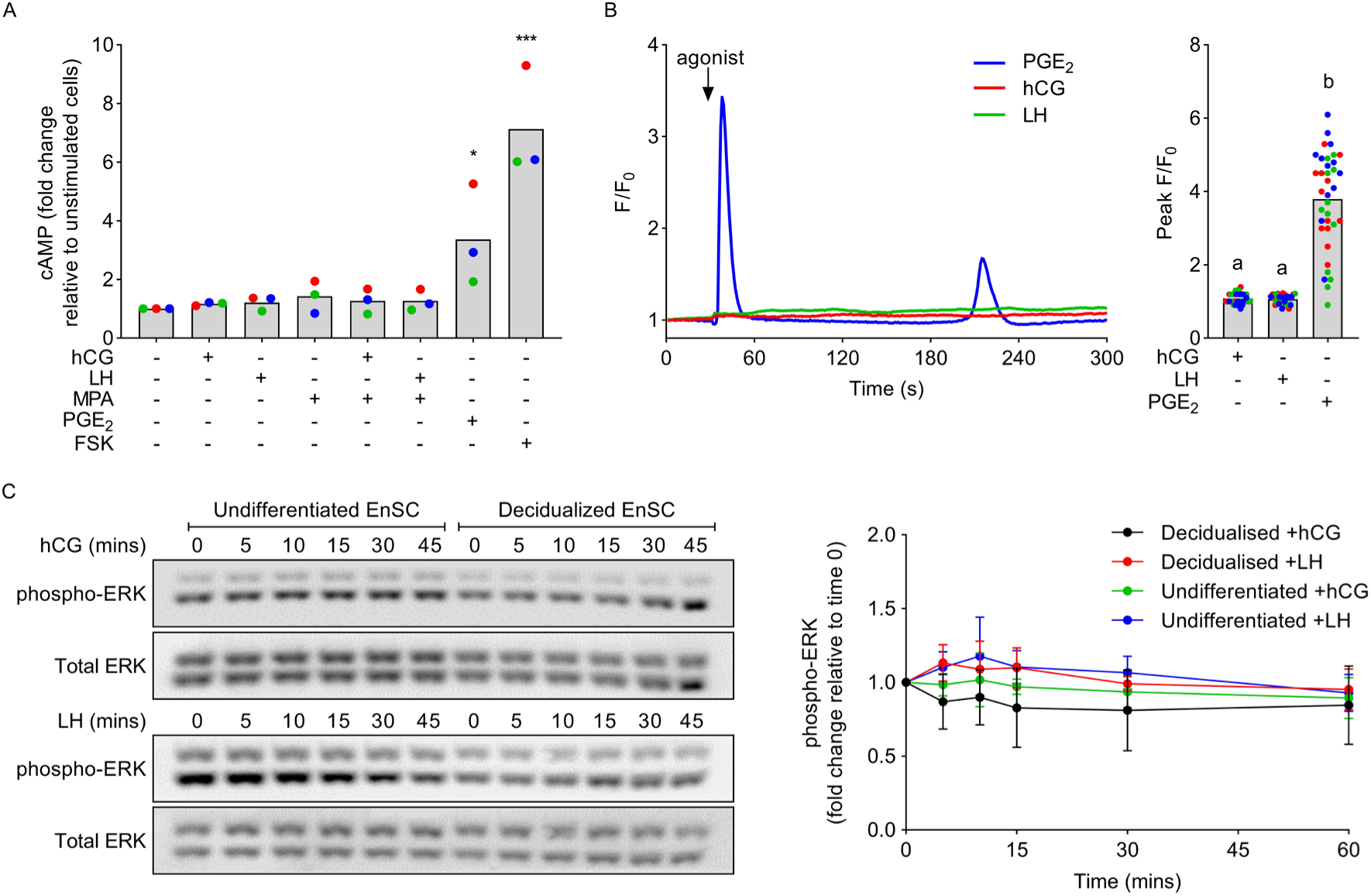
Lack of LHCGR signalling in EnSC. A) The relative change in cAMP following 5-min stimulation with 10 nM hCG or LH (with or without MPA), 2 µM PGE_2_ or 5 µM forskolin (FSK). Data are shown from 3 independent primary cultures, with bars denoting mean values. * denotes *P*<0.05 and *** *P*<0.001 (ANOVA and Dunnett’s multiple comparison test). B) Representative Ca^2+^ traces from EnSC challenged with 1 µM hCG or LH, or 2 µM PGE_2_ (left panel). Data represent one individual cell. Mean peak responses of 36 cells from 3 independent cultures (denoted as separate colours) are shown in the right panel. Different letters indicate statistical differences (*P*<0.05) between groups (ANOVA and Tukey’s multiple comparison test). C) Representative western blots showing levels of phospho-ERK in undifferentiated and decidualized EnSC following stimulation with either 10 nM hCG or LH for up to 45 minutes (left panel). Mean ± SD from 3 individual primary cultures are shown in the right panel. All changes from unstimulated cells are non-significant (ANOVA and Dunnett’s multiple comparison test).

### Lack of functional LHCGR effects in EnSC

LHCGR is known to exert proliferative and anti-apoptotic effects (1). We therefore investigated cell proliferation and viability by growing cells on microelectronic impedance sensors (xCELLigence). This permits real-time monitoring of cell cultures by quantifying changes in impedance as a representation of changes in cell shape and confluency. EnSC cultures exhibited a bi-phasic pattern of growth that stabilized into linear and sustained proliferation after 2 days. Neither the pattern, magnitude, nor rate of proliferation were affected by hCG or LH, even at high concentrations (1 µM) (Fig. 4A). Similarly, no differences in impedance were observed with hCG or LH during continuous monitoring of EnSC as they decidualized (C+M treatment) for 8 days (Fig. 4B). Furthermore, neither ligand exerted any changes in EnSC proliferation and viability after 2-days, as measured by XTT assay, even up to 10 µM (Fig. 4C).

**Figure 4.**
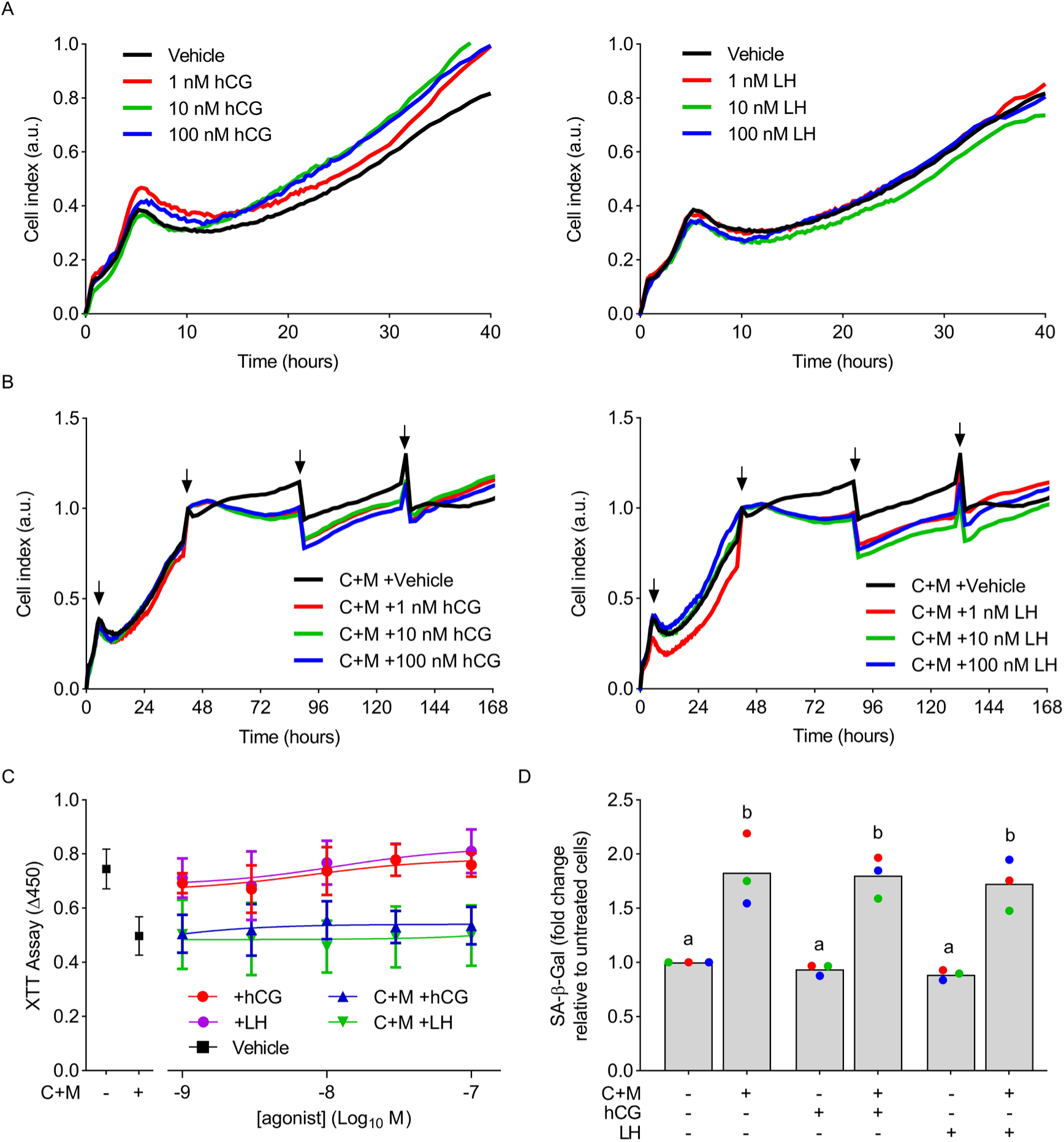
EnSC proliferation and viability are refractory to LHCGR ligands. A) xCELLigence traces showing the impact of hCG (left panels) and LH (right panels) on the proliferation of undifferentiated EnSCs. B) xCELLigence traces monitoring the effects of hCG (left panel) and LH (right panel) on decidualizing (C+M treated) EnSC over 8 days. Arrows indicate media changes. Data are representative of 3 independent primary cultures, with average data non-significant by ANOVA and Dunnett’s multiple comparison test. C) Impact of hCG and LH on cell proliferation in undifferentiated and decidualized (C+M treated) EnSC, as measured by XTT assay. Data are mean ± SD from 3 independent primary cultures. All concentrations of hCG and LH are non-significant via ANOVA and Dunnett’s multiple comparison test. D) Effects of 10 nM hCG and LH on the induction of SA-β-Gal. Data from 3 individual primary cultures are shown with different letters above bars (mean values) indicating statistical difference (*P*<0.05) from untreated cells (ANOVA and Dunnett’s multiple comparison test).

The emergence of cellular senescence in stromal cells is a key characteristic of decidualization (3, 4). We therefore investigated whether LH or hCG effected the induction of senescence-associated β-galactosidase (SA-β-Gal), a key biomarker of cellular senescence. As expected, levels of SA-β-Gal in 8 day-decidualized EnSC cultures increased 1.83 ± 0.33-fold (mean ± SD) compared to untreated controls. However, neither hCG (1.80 ± 0.19) nor LH (1.73 ± 0.23) had on any effect on SA-β-Gal activity (Fig. 4D).

Several studies propose a role for LHCGR in decidualization (refer to Table 2). Thus, we investigated whether hCG and LH regulate the induction and secretion of key decidual markers. As depicted in Fig. 5A, the production of canonical decidual markers *PRL* and *IGFBP1* co-insides with the emergence of markers that differentiate senescent and decidual cells. *IL1RL1* is a specific biomarker of decidual cells and encodes both the transmembrane (ST2L) and soluble (sST2) interleukin 1 receptor like-1, an anti-inflammatory decoy receptor that sequesters IL-33. *CLU* on the other hand codes the molecular chaperone protein clusterin (apolipoprotein J) and is enriched within the senescent subpopulation (3). First, since convergence of sustained cAMP and progesterone signalling is essential for decidualization (2), we treated EnSC continuously for 8 days with MPA in combination with 10 nM hCG or LH. In keeping with an absence of acute cAMP signalling, sustained activation of LHCGR had no impact on either the induction of decidual markers (*PRL* and *IGFBP1*) or the emergence of decidual subpopulations (*IL1RL1* and *CLU*) (Fig. 5B). To further investigate the role of LHCGR in stromal cell differentiation, EnSC cultures from 9 individual patients were decidualized with 8-br-cAMP and MPA (C+M treatment) for 8 days in combination with 10 nM hCG or LH. On average, the basal and inducted levels of decidual and subpopulation genes were refractory to hCG and LH, however considerable inter-patient variability was observed for all genes (Fig. 5C). Concomitant with transcriptional observations, we found no significant changes in secreted prolactin, IGFBP1, clusterin or sST2 when cells were decidualized for 8 days with LHCGR ligands (Fig. 5D).

**Figure 5.**
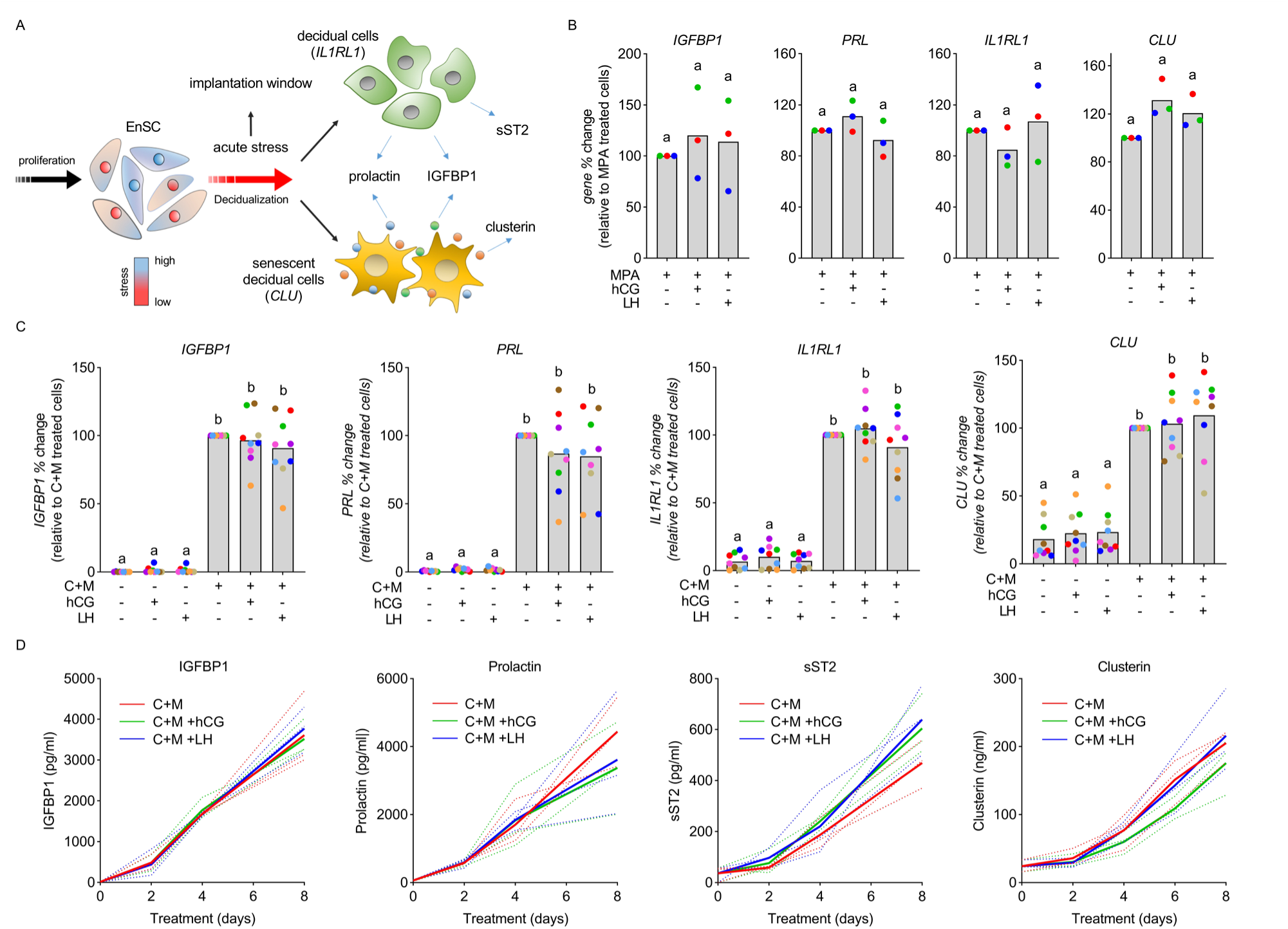
hCG and LH have no effect on EnSC decidualization. A) Schematic representation of the emergence of decidual and senescent decidual cells and the genes and secreted factors associated with each population. Modified with permission from Kong *et al* (8). B) RT-qPCR analysis of markers for decidualization (*IGFBP1* and *PRL*) and decidual (*IL1RL1*) and senescent (*CLU*) EnSC after 8-day treatment with 10 nM hCG or LH in combination with MPA, n=3. C) Impact of 10 nM hCG and LH on the decidual induction (C+M treatment) of (*IGFBP1* and *PRL*) or population specific marker genes (*IL1RL1* and *CLU*), n=9. Data are shown as individual biological replicates from independent primary cultures with bars denoting mean values. Different letters indicate statistical difference (*P*<0.05) from untreated cells (ANOVA and Dunnett’s multiple comparison test). D) Impact of 10 nM hCG and LH on the secretion of soluble decidual markers from cultured EnSC. Plots from individual patients are shown with bold lines indicating mean values. Data are non-significant via two-way ANOVA and Dunnett’s multiple comparison test.

## Discussion

hCG secretions from early embryos indirectly influence the endometrium by preventing atrophy of the corpus luteum, thus sustaining circulating progesterone and maintaining the decidua of pregnancy (2). Direct effects, however, are poorly understood. When hCG is flushed through the human and baboon uterus, endometrial changes associated with decidualisation, implantation and remodelling including upregulation of PRL, IGFBP1, leukaemia inhibitory factor (LIF), MMPs and IL-6 (46, 47) are observed. Many of these local effects have been recapitulated *in vitro* using primary endometrial cultures, and although this suggest hCG and signalling through LHCGR is important in the embryo-endometrial interface, there is contention and contradiction in the literature (summarized in Tables 1 and 2). Here, we sought to elucidate LHCGR activation within EnSC but could not detect key downstream signals or functional responses. However, despite an apparent absence of expression in the non-pregnant endometrium, we identify here, for the first time, a discrete subpopulation of stromal cells that do express LHCGR, and which may plausibly contribute to the discord between studies.

Although defining a threshold where genes can be considered functionally active poses a challenge for the analysis of transcriptomic sequencing data, there is a consensus that genes with TPM <2 are unlikely to be actively transcribed (48–50). Mean values for *LHCGR* transcripts in whole endometrial tissues were 0.15 TPM and are therefore considered non-expressed. However, insight from scRNA-seq data highlights heterogeneity in LHCGR expression, and the identification of a subpopulation of EnSC with detectable *LHCGR* transcripts. Unfortunately, *LHCGR* expressing cells did not cluster as a distinct population, and thus comparative profiling with non-expressing cells was not performed. The stability of this heterogeneity in culture as well as any inter-patient variability requires further investigation, but similar characteristics for other GPCRs in vascular and inflammatory cells have been identified (51, 52). Importantly, in these investigations, diversity in GPCR repertoire across individual cells led to subpopulation specific responses to inflammatory cues, suggesting altered pathology within individual cells. Our understanding of GPCR heterogeneity is in its infancy, and whether LHCGR positive cells are relevant in pathological conditions such as infertility, miscarriage and endometriosis remains to be seen.

The functional analysis of LHCGR is likely to be complicated further by the complex levels of regulation that govern its expression and function (53). For example, several splice variants including non-functional receptors have been identified in the human endometrium (18, 54, 55), as have immature and constitutively active receptors (10, 56, 57), plasma membrane clusters (56, 58–60), follicular stimulating hormone receptor (FSHR) heterodimers (61, 62), and receptor density-dependent signalling (63–65). Fluctuations in expression and G-protein coupling have also been linked to circadian rhythms (66). In addition, LHCGR is known to undergo rapid but transient agonist-mediated down-regulation via a LHCGR binding protein (LRBP) that accelerates the degradation of receptor mRNA (67–69), and like many other GPCRs, active LHCGR undergoes desensitisation and internalisation that not only regulates G protein signalling, but also drives alternative pathways (70, 71). The consequences of these splice variants and regulatory mechanisms as well as their sensitivity to change in culture is poorly understood, but they will profoundly impact on receptor expression and function. Although we could not identify significant effects on EnSC decidualization, we did observe inter-patient variability. For example, and consistent with previous reports (28, 32, 72), the induction of *PRL* was reduced in 3 patients when cells were decidualized with either hCG or LH. However, contradictory investigations have shown increases in *PRL* mRNA in stromal cell cultures and *in vivo* studies (33, 73). Differential responses associated with pathology have previously been identified (28) and patient-to-patient variability may be a key factor in the diversity of responses that we, and others, observe.

Whereas LH is found in most mammals, CG is an evolutionarily newer protein arising from a gene duplication event of the LHβ gene and is only found in primates (74, 75). Compared to LH, hCG has more carbohydrate moieties on the β subunit resulting in several glycoforms including a hyperglycosylated variant that predominates in early pregnancy (1, 76–78). Differences between urinary and recombinantly-sourced hCG and the potency of the hyperglycosylated variant may contribute to the range of responses reported (79, 80). This is complicated further by non-LHCGR mediated effects through mannose and other lectin-like receptors. These receptors recognize terminal mannose, N-GlcNAc and fructose residues on glycans attached to proteins and are known to interact with pituitary hormones include TSH and LH (42–44). They are abundant in a range of immune cells, many of which, including uNK cells, macrophages, dendritic cells and regulatory T cells, infiltrate the maternal decidua. In this study, hCG augmented uNK cell proliferation. A response previously linked to activation of mannose receptors (44). There is also further evidence that hCG stimulates cytokine release from peripheral blood mononuclear cells (PBMC) including IL-18, IL-1β, IL-6, and TNF-α, interferon γ and soluble IL-2 receptor (42, 81–83). In many cases, responses were sensitive to excess D-mannose and refractory to deglycosylated hCG, emphasising the importance of glyco-conjugates. Given the critical function of uNK cells and cytokine release during decidualisation and implantation (3, 4), immune cell/lectin receptor activation cannot be excluded from the endometrial responses to uterine hCG flushing. Here, we showed that both stromal and immune cells predominantly express the type 2 mannose receptor (*MRC2*), whereas *LHCGR* was absent in immune cells. Consequently, the roles of hCG within the endometrium must be evaluated within immune cells and via alternative receptors as well as canonical LHCGR activation. It is therefore necessary for researchers to design future experiments that deplete cells of LHCGR and mannose receptors to identify receptor activation.

From an evolutionary context, the importance of hCG at the endometrial-embryo interface during early pregnancy is undermined by certain characteristics which no longer make it an honest signal of embryo fitness (84). Considering the excessive concentrations of hCG observed in early pregnancy, the multiple glycoforms and gene duplications and its extended half-life, it makes little sense to rely solely on hCG as a gauge for embryo quality. Thus, the role of hCG-LHCGR within the endometrium, at least from the evolutionary perspective of embryo selection, loses importance. However, the endometrium does undergo profound adaptations during pregnancy, and these adaptations can be intrinsically modified in response to embryonic and environmental cues (2). How the activation and regulation of endometrial LHCGR is modified during pregnancy, especially in response to embryonic hCG, is unknown, and highlights the need for models that recapitulate the multicellular endometrial environment during early pregnancy.

In summary, we report that transcripts for LHCGR were largely absent in non-pregnant endometrial tissues and EnSC cultures but were identified in a small subpopulation of stromal cells. Despite confirming bioactivity in HEK-LHCGR cells, neither hCG nor LH generated key downstream signals or functional responses in EnSC, and we oppose any assertion that LHCGR is functionally active in endometrial stromal cells. However, we highlight the need for caution when interpreting experimental data that ignores both heterogeneity within endometrial cell populations and non-canonical hCG/LH-LHCGR signalling through C-type lectin and mannose receptors.

## Author Contributions

Conceptualization, P.J.B., A.H and J.J.B; Methodology, P.J.B., E.S.L and C.S.K; Investigation, P.J.B., E.S.L., O.M and C.S.K; Writing P.J.B., A.H., O.M and J.J.B; Funding Acquisition, A.H and J.J.B; Resources, J.J.B; Supervision, P.J.B., A.H and J.J.B.

## Acknowledgements

The authors are indebted to all the women who participated in this study and generously donated endometrial biopsies for research. We are also grateful to those who facilitated their participation including Professor Siobhan Quenby, Dr Joshua Odendaal and Dr Amelia Hawkes. This work was supported by a UKRI BBSRC Award to Dr Aylin Hanyaloglu (BB/S001565/1) and by funds from the Tommy’s National Miscarriage Research Centre and Wellcome Trust Investigator Award to Professor Jan Brosens (212233/Z/18/Z).

## Disclaimer

The authors have no conflicts of interest to declare.

## Supplementary Information

**Figure S1.**
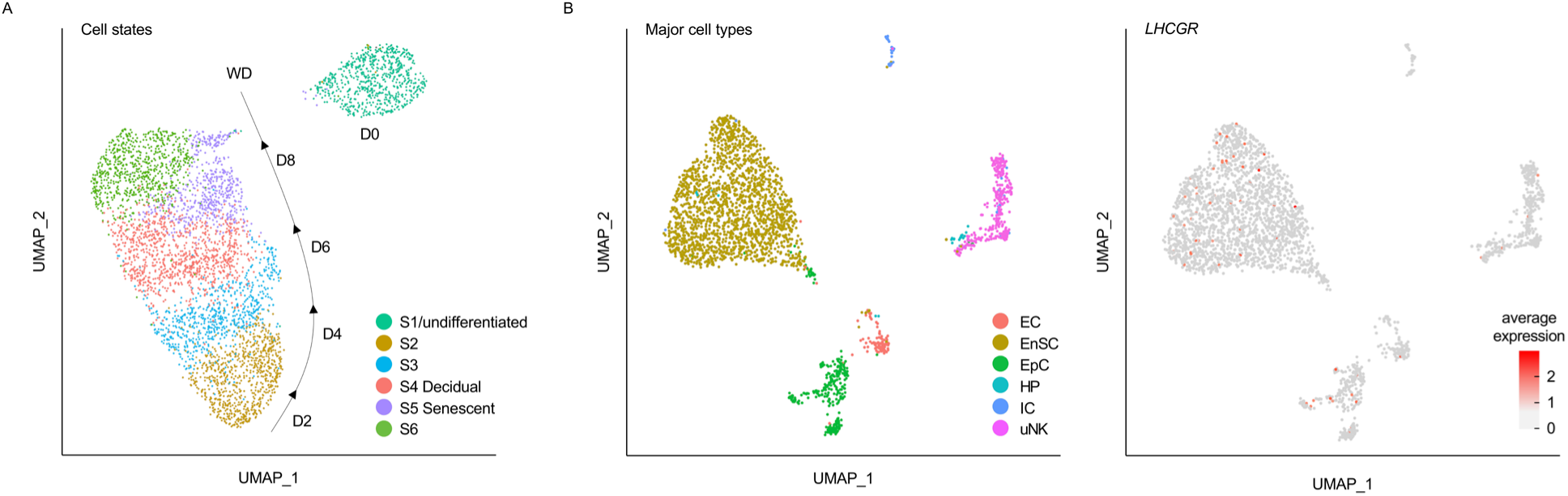
Transcriptional clustering of endometrial stromal cells. UMAP (uniform manifold approximation and projection) projections from scRNA-seq analysis of undifferentiated (day 0) EnSC and those treated with C+M for up to 8 days before withdrawal (WD) (GEO GSE127918). Cells are colour coded according to transcriptional states (S1-6). B) UMAP projections from scRNA-seq data of endometrial biopsies (GEO GSE127918), identifying major endometrial cell types (endothelial (EC), stromal cells (EnSC), epithelial cells (EpC), highly proliferative (HP), immune (IC) and uterine natural killer (uNK) cells). Data adapted from Lucas *et al*, 2020 with permission (3). C) UMAP projections from the same data set depicting *LHCGR* expression across cell types.

**Figure S2.**
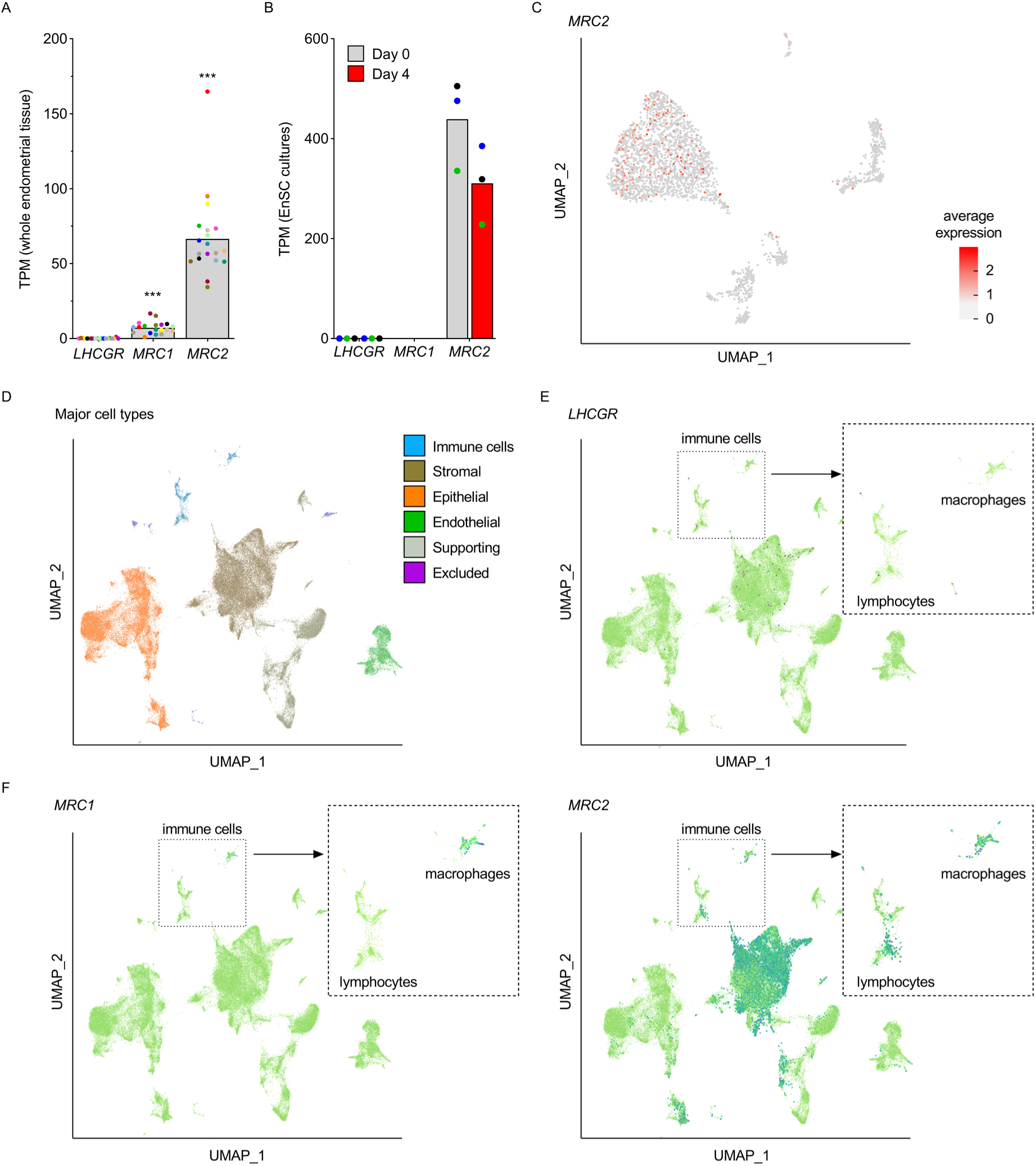
Endometrial mannose receptor expression. A) Comparison of *LHCGR* expression with the mannose receptor C-type 1 (*MRC1*) and 2 (*MRC2*) in bulk RNA-seq data from whole endometrial tissues (GEO GSE65102). Data points from individual patients are colour matched and shown together with bars denoting mean, n= 20. *** denotes *P*<0.001 by ANOVA and Dunnett’s multiple comparison using *LHCGR* as the comparator. B) Expression of *LHCGR*, *MRC1* and *MRC2* in bulk RNA-seq data from undifferentiated EnSC (day 0) and cells decidualized with 8-br-cAMP and MPA (C+M) for 4 days (GEO GSE104721). C) UMAP projections depicting *MRC2* expression across different endometrial cells (GEO GSE127918). Refer to Fig. S1B for cell types. *MRC1* was not expressed. D) UMAP projections from scRNA-seq data identifying major cell types within whole endometrial biopsies (37). E) UMAP projection depicting *LHCGR* expression across endometrial cells identified in D. F) UMPA projections depicting *MRC1* (F, left panel) and *MRC2* (F, right panel) expression within the same dataset. Inserts highlight the expression of *LHCGR*, *MRC1* and *MRC2* within immune cells.

**Table S1.** Patient demographics

| **Endometrial biopsies used for EnSC cultures** |  |  |  |  |
| --- | --- | --- | --- | --- |
| **Figure** | **(n)** | **Age** | **BMI** | **LH+** |
| **2E** | 13 | 36 (33.75-37) | 23.5 (19-28.25) | 9 (8-10) |
| **3A** | 3 | 39 (37-39.5) | 26 (20-26.25) | 8 (7-8) |
| **3B** | 3 | 38 (33-39.5) | 21 (20-23.5) | 7 (5-8) |
| **3C** | 3 | 36 (30-39) | 23 (22-23) | 11 (8-11) |
| **4A-B** | 3 | 35 (35-38) | 23 (22-23.75) | 9 (8-9.5) |
| **4C-D** | 3 | 39 (37-39.5) | 26 (20-26.25) | 8 (7-8) |
| **5B and D** | 3 | 35 (34-35.5) | 25 (25-26) | 8 (7-8.5) |
| **5C** | 9 | 38 (34-41) | 25 (22-27) | 8 (7-9) |

All data are median (interquartile range, Q1-Q3).

## References

1. Casarini, L., Santi, D., Brigante, G., and Simoni, M. (2018) Two Hormones for One Receptor: Evolution, Biochemistry, Actions, and Pathophysiology of LH and hCG. Endocrine Reviews 39, 549–592

2. Gellersen, B., and Brosens, J. J. (2014) Cyclic Decidualization of the Human Endometrium in Reproductive Health and Failure. Endocrine Reviews 35, 851–905

3. Lucas, E. S., Vrljicak, P., Muter, J., Diniz-da-Costa, M. M., Brighton, P. J., Kong, C.-S., Lipecki, J., Fishwick, K. J., Odendaal, J., Ewington, L. J., Quenby, S., Ott, S., and Brosens, J. J. (2020) Recurrent pregnancy loss is associated with a pro-senescent decidual response during the peri-implantation window. Communications Biology 3, 37

4. Brighton, P. J., Maruyama, Y., Fishwick, K., Vrljicak, P., Tewary, S., Fujihara, R., Muter, J., Lucas, E. S., Yamada, T., Woods, L., Lucciola, R., Lee, Y. H., Takeda, S., Ott, S., Hemberger, M., Quenby, S., and Brosens, J. J. (2017) Clearance of senescent decidual cells by uterine natural killer cells in cycling human endometrium. eLife 6

5. Critchley, H. O. D., Maybin, J. A., Armstrong, G. M., and Williams, A. R. W. (2020) Physiology of the endometrium and regulation of menstruation. Physiological Reviews 100, 1149–1179

6. Ewington, L. J., Tewary, S., and Brosens, J. J. (2019) New insights into the mechanisms underlying recurrent pregnancy loss. Journal of Obstetrics and Gynaecology Research 45, 258–265

7. Haig, D. (2019) Cooperation and conflict in human pregnancy. Current Biology 29, R455–R458

8. Kong, C. S., Ordonez, A. A., Turner, S., Tremaine, T., Muter, J., Lucas, E. S., Salisbury, E., Vassena, R., Tiscornia, G., Fouladi-Nashta, A. A., Hartshorne, G., Brosens, J. J., and Brighton, P. J. (2021) Embryo biosensing by uterine natural killer cells determines endometrial fate decisions at implantation. Faseb Journal 35, e21336

9. Brosens, J. J., Salker, M. S., Teklenburg, G., Nautiyal, J., Salter, S., Lucas, E. S., Steel, J. H., Christian, M., Chan, Y. W., Boomsma, C. M., Moore, J. D., Hartshorne, G. M., Sucurovic, S., Mulac-Jericevic, B., Heijnen, C. J., Quenby, S., Koerkamp, M. J. G., Holstege, F. C. P., Shmygol, A., and Macklon, N. S. (2014) Uterine Selection of Human Embryos at Implantation. Scientific Reports 4

10. Ascoli, M., Fanelli, F., and Segaloff, D. L. (2002) The lutropin/choriocrctnadotropin receptor, a 2002 perspective. Endocrine Reviews 23, 141–174

11. Xie, Y. B., Wang, H. Y., and Segaloff, D. L. (1990) Extracellular domain of lutropin choriogonadotropin receptor expressed in transfeced cells binds choriogonadotropin with high-affinity. Journal of Biological Chemistry 265, 21411–21414

12. Choi, J., and Smitz, J. (2014) Luteinizing hormone and human chorionic gonadotropin: Origins of difference. Molecular and Cellular Endocrinology 383, 203–213

13. Breen, S. M., Andric, N., Ping, T., Xie, F., Offermans, S., Gossen, J. A., and Ascoli, M. (2013) Ovulation Involves the Luteinizing Hormone-Dependent Activation of G(q/11) in Granulosa Cells. Molecular Endocrinology 27, 1483–1491

14. Gilchrist, R. L., Ryu, K. S., Ji, I. H., and Ji, T. H. (1996) The luteinizing hormone chorionic gonadotropin receptor has distinct transmembrane conductors for cAMP and inositol phosphate signals. Journal of Biological Chemistry 271, 19283–19287

15. Reshef, E., Lei, Z. M., Rao, C. V., Pridham, D. D., Chegini, N., and Luborsky, J. L. (1990) The presence of gonadotropin receptors in nonpregnant human uterus, human placenta, fetal membranes, and decidua. J. Clin. Endocrinol. Metab. 70, 421–430

16. Lei, Z. M., Toth, P., Rao, C. V., and Pridham, D. (1993) Novel co-expression of human chorionic-gonadotropin (hCG) human luteinizing-hormone receptors and their ligand hCG in human fallopian-tubes. J. Clin. Endocrinol. Metab. 77, 863–872

17. Zuo, J., Lei, Z. M., and Rao, C. V. (1994) Human myometrial chorionic-gonadotropin luteinizing-hormone receptors in preterm and term deliveries. J. Clin. Endocrinol. Metab. 79, 907–911

18. Stewart, E. A., Sahakian, M., Rhoades, A., Van Voorhis, B. J., and Nowak, R. A. (1999) Messenger ribonucleic acid for the gonadal luteinizing hormone human chorionic gonadotropin receptor is not present in human endometrium. Fertility and Sterility 71, 368–372

19. Pakarainen, T., Ahtiainen, P., Zhang, F.-P., Rulli, S., Poutanen, M., and Huhtaniemi, I. (2007) Extragonadal LH/hCG action—Not yet time to rewrite textbooks. Molecular and Cellular Endocrinology 269, 9–16

20. Licht, P., von Wolff, M., Berkholz, A., and Wildt, L. (2003) Evidence for cycle-dependent expression of full-length human chorionic gonadotropin/luteinizing hormone receptor mRNA in human endometrium and decidua. Fertility and Sterility 79, 718–723

21. Srisuparp, S., Strakova, Z., Brudney, A., Mukherjee, S., Reierstad, S., Hunzicker-Dunn, M., and Fazleabas, A. T. (2003) Signal transduction pathways activated by chorionic gonadotropin in the primate endometrial epithelial cells. Biology of Reproduction 68, 457–464

22. Perrier d’Hauterive, S., Charlet-Renard, C., Berndt, S., Dubois, M., Munaut, C., Goffin, F., Hagelstein, M. T., Noel, A., Hazout, A., Foidart, J. M., and Geenen, V. (2004) Human chorionic gonadotropin and growth factors at the embryonic-endometrial interface control leukemia inhibitory factor (LIF) and interleukin 6 (IL-6) secretion by human endometrial epithelium. Human Reproduction 19, 2633–2643

23. Banerjee, P., Sapru, K., Strakova, Z., and Fazleabas, A. T. (2009) Chorionic Gonadotropin Regulates Prostaglandin E Synthase via a Phosphatidylinositol 3-Kinase-Extracellular Regulatory Kinase Pathway in a Human Endometrial Epithelial Cell Line: Implications for Endometrial Responses for Embryo Implantation. Endocrinology 150, 4326–4337

24. Evans, J., and Salamonsen, L. A. (2013) Too much of a good thing? Experimental evidence suggests prolonged exposure to hCG is detrimental to endometrial receptivity. Human Reproduction 28, 1610–1619

25. Bernardini, L., Moretti-Rojas, I., Brush, M., Rojas, F., and Balmaceda, J. (2013) Failure of hCG/LH receptors to stimulate the transmembrane effector adenylyl cyclase in human endometrium. Advances in Bioscience and Biotechnology 4, 949

26. Tapia-Pizarro, A., Archiles, S., Argandona, F., Valencia, C., Zavaleta, K., Johnson, M. C., Gonzalez-Ramos, R., and Devoto, L. (2017) hCG activates Epac-Erk1/2 signaling regulating Progesterone Receptor expression and function in human endometrial stromal cells. Molecular Human Reproduction 23, 393–405

27. Sacchi, S., Sena, P., Degli Esposti, C., Lui, J., and La Marca, A. (2018) Evidence for expression and functionality of FSH and LH/hCG receptors in human endometrium. J. Assist. Reprod. Genet. 35, 1703–1712

28. Salker, M., Teklenburg, G., Molokhia, M., Lavery, S., Trew, G., Aojanepong, T., Mardon, H. J., Lokugamage, A. U., Rai, R., Landles, C., Roelen, B. A. J., Quenby, S., Kuijk, E. W., Kavelaars, A., Heijnen, C. J., Regan, L., Macklon, N. S., and Brosens, J. J. (2010) Natural Selection of Human Embryos: Impaired Decidualization of Endometrium Disables Embryo-Maternal Interactions and Causes Recurrent Pregnancy Loss. Plos One 5, e10287

29. West, C., Hanyaloglu, A., Brosens, J., Lavery, S., and Panay, N. (2014) Regulation of the LH/CG receptor signalling in human endometrium. Endorcine Abstracts 34, P332

30. Tang, B. Q., and Gurpide, E. (1993) Direct effect of gonadotropins on decidualization of human endometrial stromal cells. J. Steroid Biochem. Mol. Biol. 47, 115–121

31. Chatterjee, A., Jana, N. R., and Bhattacharya, S. (1997) Stimulation of cyclic AMP, 17 beta-oestradiol and protein synthesis by human chorionic gonadotrophin in human endometrial cells. Human Reproduction 12, 1903–1908

32. Kasahara, K., Takakura, K., Takebayashi, K., Kimura, F., Nakanishi, K., and Noda, Y. (2001) The role of human chorionic gonadotropin on decidualization of endometrial stromal cells in vitro. J. Clin. Endocrinol. Metab. 86, 1281–1286

33. Han, S. W., Lei, Z. M., and Rao, C. V. (1999) Treatment of human endometrial stromal cells with chorionic gonadotropin promotes their morphological and functional differentiation into decidua. Molecular and Cellular Endocrinology 147, 7–16

34. Kajihara, T., Uchino, S., Suzuki, M., Itakura, A., Brosens, J. J., and Ishihara, O. (2011) Human chorionic gonadotropin confers resistance to oxidative stress-induced apoptosis in decidualizing human endometrial stromal cells. Fertility and Sterility 95, 1302–1307

35. Lucas, E. S., Dyer, N. P., Murakami, K., Lee, Y. H., Chan, Y. W., Grimaldi, G., Muter, J., Brighton, P. J., Moore, J. D., Patel, G., Chan, J. K. Y., Takeda, S., Lam, E. W. F., Quenby, S., Ott, S., and Brosens, J. J. (2016) Loss of Endometrial Plasticity in Recurrent Pregnancy Loss. Stem Cells 34, 346–356

36. Muter, J., Alam, M. T., Vrljicak, P., Barros, F. S. V., Ruane, P. T., Ewington, L. J., Aplin, J. D., Westwood, M., and Brosens, J. J. (2018) The Glycosyltransferase EOGT Regulates Adropin Expression in Decidualizing Human Endometrium. Endocrinology 159, 994–1004

37. Garcia-Alonso, L., Handfield, L. F., Roberts, K., Nikolakopoulou, K., Fernando, R. C., Gardner, L., Woodhams, B., Arutyunyan, A., Polanski, K., Hoo, R., Sancho-Serra, C., Li, T., Kwakwa, K., Tuck, E., Lorenzi, V., Massalha, H., Prete, M., Kleshchevnikov, V., Tarkowska, A., Porter, T., Mazzeo, C. I., van Dongen, S., Dabrowska, M., Vaskivskyi, V., Mahbubani, K. T., Park, J. E., Jimenez-Linan, M., Campos, L., Kiselev, V. Y., Lindskog, C., Ayuk, P., Prigmore, E., Stratton, M. R., Saeb-Parsy, K., Moffett, A., Moore, L., Bayraktar, O. A., Teichmann, S. A., Turco, M. Y., and Vento-Tormo, R. (2021) Mapping the temporal and spatial dynamics of the human endometrium in vivo and in vitro. Nature Genetics 53, 1698

38. Megill, C., Martin, B., Weaver, C., Bell, S., Prins, L., Badajoz, S., McCandless, B., Pisco, A. O., Kinsella, M., Griffin, F., Kiggins, J., Haliburton, G., Mani, A., Weiden, M., Dunitz, M., Lombardo, M., Huang, T., Smith, T., Chambers, S., Freeman, J., Cool, J., and Carr, A. (2021) cellxgene: a performant, scalable exploration platform for high dimensional sparse matrices. bioRxiv, 438318

39. Barros, F., Brosens, J., and Brighton, P. (2016) Isolation and Primary Culture of Various Cell Types from Whole Human Endometrial Biopsies. Bio-Protocol 6, e2028

40. Jean-Alphonse, F., Bowersox, S., Chen, S., Beard, G., Puthenveedu, M. A., and Hanyaloglu, A. C. (2014) Spatially Restricted G Protein-coupled Receptor Activity via Divergent Endocytic Compartments. Journal of Biological Chemistry 289, 3960–3977

41. Macosko, E. Z., Basu, A., Satija, R., Nemesh, J., Shekhar, K., Goldman, M., Tirosh, I., Bialas, A. R., Kamitaki, N., Martersteck, E. M., Trombetta, J. J., Weitz, D. A., Sanes, J. R., Shalek, A. K., Regev, A., and McCarroll, S. A. (2015) Highly Parallel Genome-wide Expression Profiling of Individual Cells Using Nanoliter Droplets. Cell 161, 1202–1214

42. Kosaka, K., Fujiwara, H., Tatsumi, K., Yoshioka, S., Sato, Y., Egawa, H., Higuchi, T., Nakayama, T., Ueda, M., Maeda, M., and Fujii, S. (2002) Human chorionic gonadotropin (HCG) activates monocytes to produce interleukin-8 via a different pathway from luteinizing hormone/HCG receptor system. Journal of Clinical Endocrinology & Metabolism 87, 5199–5208

43. Simpson, D. Z., Hitchen, P. G., Elmhirst, E. L., and Taylor, M. E. (1999) Multiple interactions between pituitary hormones and the mannose receptor. Biochemical Journal 343, 403–411

44. Kane, N., Kelly, R., Saunders, P. T. K., and Critchley, H. O. D. (2009) Proliferation of Uterine Natural Killer Cells Is Induced by Human Chorionic Gonadotropin and Mediated via the Mannose Receptor. Endocrinology 150, 2882–2888

45. Mi, Y. L., Lin, A., Fiete, D., Steirer, L., and Baenziger, J. U. (2014) Modulation of Mannose and Asialoglycoprotein Receptor Expression Determines Glycoprotein Hormone Half-life at Critical Points in the Reproductive Cycle. Journal of Biological Chemistry 289, 12157–12167

46. Licht, P., Russu, V., and Wildt, L. (2001) On the role of human chorionic gonadotropin (hCC) in the embryo-endometrial microenvironment: Implications for differentiation and implantation. Seminars in Reproductive Medicine 19, 37–47

47. Sherwin, J. R. A., Sharkey, A. M., Cameo, P., Mavrogianis, P. M., Catalano, R. D., Edassery, S., and Fazleabas, A. T. (2007) Identification of novel genes regulated by chorionic gonadotropin in baboon endometrium during the window of implantation. Endocrinology 148, 618–626

48. Mika, K., Marinic, M., Singh, M., Muter, J., Brosens, J. J., and Lynch, V. J. (2021) Evolutionary transcriptomics implicates new genes and pathways in human pregnancy and adverse pregnacy outcomes. eLIFE, 10:e69584

49. Wagner, G. P., Kin, K., and Lynch, V. J. (2012) Measurement of mRNA abundance using RNA-seq data: RPKM measure is inconsistent among samples. Theory in Biosciences 131, 281–285

50. Wagner, G. P., Kin, K., and Lynch, V. J. (2013) A model based criterion for gene expression calls using RNA-seq data. Theory in Biosciences 132, 159–164

51. Kaur, H., Carvalho, J., Looso, M., Singh, P., Chennupati, R., Preussner, J., Guenther, S., Albarran-Juarez, J., Tischner, D., Classen, S., Offermanns, S., and Wettschureck, N. (2019) Single-cell profiling reveals heterogeneity and functional patterning of GPCR expression in the vascular system. Nature Communications 10, 15700

52. Tischner, D., Grimm, M., Kaur, H., Staudenraus, D., Carvalho, J., Looso, M., Guenther, S., Wanke, F., Moos, S., Siller, N., Breuer, J., Schwab, N., Zipp, F., Waisman, A., Kurschus, F. C., Offermanns, S., and Wettschureck, N. (2017) Single-cell profiling reveals GPCR heterogeneity and functional patterning during neuroinflammation. JCI Insight 2, e95063

53. Menon, K. M. J., Munshi, U. M., Clouser, C. L., and Nair, A. K. (2004) Regulation of luteinizing hormone/human chorionic gonadotropin receptor expression: A perspective. Biology of Reproduction 70, 861–866

54. Minegishi, T., Tano, M., Abe, Y., Nakamura, K., Ibuki, Y., and Miyamoto, K. (1997) Expression of luteinizing hormone human chorionic gonadotrophin (LH/HCG) receptor mRNA in the human ovary. Molecular Human Reproduction 3, 101–107

55. Madhra, M., Gay, E., Fraser, H. M., and Duncan, W. C. (2004) Alternative splicing of the human luteal LH receptor during luteolysis and maternal recognition of pregnancy. Molecular Human Reproduction 10, 599–603

56. Tao, Y. X., Johnson, N. B., and Segaloff, D. L. (2004) Constitutive and agonist-dependent self-association of the cell surface human lutropin receptor. Journal of Biological Chemistry 279, 5904–5914

57. Lei, Y., Hagen, G. M., Smith, S. M. L., Liu, J. I., Barisas, G., and Roess, D. A. (2007) Constitutively-active human LH receptors are self-associated and located in rafts. Molecular and Cellular Endocrinology 260, 65–72

58. Roess, D. A., Horvat, R. D., Munnelly, H., and Barisas, B. G. (2000) Luteinizing hormone receptors are self-associated in the plasma membrane. Endocrinology 141, 4518–4523

59. Horvat, R. D., and Roess, D. A. (2000) Desensitized LH receptors are self-associated in large, slowly diffusing complexes. Biology of Reproduction 62, 106–106

60. Smith, S. M. L., Lei, Y., Liu, J. J., Cahill, M. E., Hagen, G. M., Barisas, B. G., and Roess, D. A. (2006) Luteinizing hormone receptors translocate to plasma membrane microdomains after binding of human chorionic gonadotropin. Endocrinology 147, 1789–1795

61. Zhang, M. L., Feng, X. Y., Guan, R. B., Hebert, T. E., and Segaloff, D. L. (2009) A cell surface inactive mutant of the human lutropin receptor (hLHR) attenuates signaling of wild-type or constitutively active receptors via heterodimerization. Cell. Signal. 21, 1663–1671

62. Feng, X. Y., Zhang, M. L., Guan, R. B., and Segaloff, D. L. (2013) Heterodimerization Between the Lutropin and Follitropin Receptors is Associated With an Attenuation of Hormone-Dependent Signaling. Endocrinology 154, 3925–3930

63. Zhu, X., Gilbert, S., Birnbaumer, M., and Birnbaumer, L. (1994) Dual signaling potential is common among g(s)-coupled receptors and dependent on receptor density. Mol. Pharmacol. 46, 460–469

64. Donadeu, F. X., and Ascoli, M. (2005) The differential effects of the gonadotropin receptors on aromatase expression in primary cultures of immature rat granulosa cells are highly dependent on the density of receptors expressed and the activation of the inositol phosphate cascade. Endocrinology 146, 3907–3916

65. Casarini, L., Reiter, E., and Simoni, M. (2016) beta-arrestins regulate gonadotropin receptor-mediated cell proliferation and apoptosis by controlling different FSHR or LHCGR intracellular signaling in the hGL5 cell line. Molecular and Cellular Endocrinology 437, 11–21

66. Gilioli, L., Marino, M., Simon, M., and Cassarini, L. (2017) The regulation of LHCGR-dependent signaling is linked to circadian gene expression. Endocrine Abstracts 49, EP1131

67. Kishi, H., Kitahara, Y., Imai, F., Nakao, K., and Suwa, H. (2018) Expression of the gonadotropin receptors during follicular development. Reproductive Medicine and Biology 17, 11–19

68. Nair, A. K., Kash, J. C., Peegel, H., and Menon, K. M. J. (2002) Post-transcriptional regulation of luteinizing hormone receptor mRNA in the ovary by a novel mRNA-binding protein. Journal of Biological Chemistry 277, 21468–21473

69. Kash, J. C., and Menon, K. M. J. (1998) Identification of a hormonally regulated luteinizing hormone human chorionic gonadotropin receptor mRNA binding protein - Increased mRNA binding during receptor down-regulation. Journal of Biological Chemistry 273, 10658–10664

70. Sposini, S., and Hanyaloglu, A. C. (2018) Driving gonadotrophin hormone receptor signalling: the role of membrane trafficking. Reproduction 156, R195–R208

71. Johnson, G. P., and Jonas, K. C. (2019) Mechanistic insight into how gonadotropin hormone receptor complexes direct signaling†. Biology of Reproduction 102, 773–783

72. Fluhr, H., Krenzer, S., Deperschmidt, M., Zwirner, M., Wallwiener, D., and Licht, P. (2006) Human chorionic gonadotropin inhibits insulin-like growth factor-binding protein-1 and prolactin in decidualized human endometrial stromal cells. Fertility and Sterility 86, 236–238

73. Licht, P., Losch, A., Dittrich, R., Neuwinger, J., Siebzehnrubl, E., and Wildt, L. (1998) Novel insights into human endometrial paracrinology and embryo-maternal communication by intrauterine microdialysis. Human Reproduction Update 4, 532–538

74. Fiddes, J. C., and Goodman, H. M. (1980) The cDNA for the beta-subunit of human chorionic-gonadotropin suggests evolution of a gene by readthrough into the 3’-untranslated region. Nature 286, 684–687

75. Maston, G. A., and Ruvolo, M. (2002) Chorionic gonadotropin has a recent origin within primates and an evolutionary history of selection. Mol. Biol. Evol. 19, 320–335

76. Cole, L. A. (2012) hCG, five independent molecules. Clin. Chim. Acta 413, 48–65

77. Kovalevskaya, G., Birken, S., Kakuma, T., and O’Connor, J. F. (1999) Early pregnancy human chorionic gonadotropin (hCG) isoforms measured by an immunometric assay for choriocarcinoma-like hCG. J. Endocrinol. 161, 99–106

78. Stenman, U. H., Birken, S., Lempiainen, A., Hotakainen, K., and Alfthan, H. (2011) Elimination of Complement Interference in Immunoassay of Hyperglycosylated Human Chorionic Gonadotropin. Clinical Chemistry 57, 1075–1077

79. Koistinen, H., Koel, M., Peters, M., Rinken, A., Lundin, K., Tuuri, T., Tapanainen, J. S., Alfthan, H., Salumets, A., Stenman, U. H., and Lavogina, D. (2019) Hyperglycosylated hCG activates LH/hCG-receptor with lower activity than hCG. Molecular and Cellular Endocrinology 479, 103–109

80. Kajihara, T., Tochigi, H., Uchino, S., Itakura, A., Brosens, J. J., and Ishihara, O. (2011) Differential effects of urinary and recombinant chorionic gonadotropin on oxidative stress responses in decidualizing human endometrial stromal cells. Placenta 32, 592–597

81. Schafer, A., Pauli, G., Friedmann, W., and Dudenhausen, J. W. (1992) Human choriogonadotropin (hCG) and placental-lactogen (HPL) inhibit interleukin-2 (IL-2) and increase interleukin-1-beta (IL-1-beta), interleukin-6 (IL-6) and tumor-necrosis-factor (TNF-alpha) expression in monocyte cell-cultures. Journal of Perinatal Medicine 20, 233–240

82. Yousefi, S., Karamlou, K., Vaziri, N., Carandang, G., Ocariz, J., and Cesario, T. (1993) The effect of gonadotropins on the production of human interferon-gamma by mononuclear-cells. Journal of Interferon Research 13, 213–220

83. Komorowski, J., Gradowski, G., and Stepien, H. (1997) Effects of hCG and beta-hCG on IL-2 and sIL-2R secretion from human peripheral blood mononuclear cells: A dose-response study in vitro. Immunology Letters 59, 29–33

84. McCoy, D. E., and Haig, D. (2020) Embryo Selection and Mate Choice: Can ‘Honest Signals’ Be Trusted? Trends in Ecology & Evolution 35, 308–318

85. Han, S. W., Lei, Z. M., and Rao, C. V. (1997) Homologous down-regulation of luteinizing hormone chorionic gonadotropin receptors by increasing the degradation of receptor transcripts in human uterine endometrial stromal cells. Biology of Reproduction 57, 158–164

86. Han, S. W., Lei, Z. M., and Rao, C. V. (1996) Up-regulation of cyclooxygenase-2 gene expression by chorionic gonadotropin during the differentiation of human endometrial stromal cells into decidua. Endocrinology 137, 1791–1797

87. Berndt, S., d’Hauterive, S. P., Blacher, S., Pequeux, C., Lorquet, S., Munaut, C., Applanat, M., Herve, M. A., Lamande, N., Corvol, P., van den Brule, F., Frankenne, F., Poutanen, M., Huhtaniemi, I., Geenen, V., Noel, A., and Foidart, J. M. (2006) Angiogenic activity of human chorionic gonadotropin through LH receptor activation on endothelial and epithelial cells of the endometrium. Faseb Journal 20, 2630

88. Paiva, P., Hannan, N. J., Hincks, C., Meehan, K. L., Pruysers, E., Dimitriadis, E., and Salamonsen, L. A. (2011) Human chorionic gonadotrophin regulates FGF2 and other cytokines produced by human endometrial epithelial cells, providing a mechanism for enhancing endometrial receptivity. Human Reproduction 26, 1153–1162

